# A pressure-jump EPR system to monitor millisecond conformational exchange rates of spin-labeled proteins

**DOI:** 10.1101/2024.05.07.593074

**Authors:** Julian D. Grosskopf, Jason W. Sidabras, Christian Altenbach, Jim R. Anderson, Richard R. Mett, Robert A. Strangeway, James S. Hyde, Wayne L. Hubbell, Michael T. Lerch

## Abstract

Site-directed spin labeling electron paramagnetic resonance (SDSL-EPR) using nitroxide spin labels is a well-established technology for mapping site-specific secondary and tertiary structure and for monitoring conformational changes in proteins of any degree of complexity, including membrane proteins, with high sensitivity. SDSL-EPR also provides information on protein dynamics in the time scale of ps-μs using continuous wave lineshape analysis and spin lattice relaxation time methods. However, the functionally important time domain of μs-ms, corresponding to large-scale protein motions, is inaccessible to those methods. To extend SDSL-EPR to the longer time domain, the perturbation method of pressure-jump relaxation is implemented. Here, we describe a complete high-pressure EPR system at Q-band for both static pressure and millisecond-timescale pressure-jump measurements on spin-labeled proteins. The instrument enables pressure jumps both up and down from any holding pressure, ranging from atmospheric pressure to the maximum pressure capacity of the system components (∼3500 bar). To demonstrate the utility of the system, we characterize a local folding-unfolding equilibrium of T4 lysozyme. The results illustrate the ability of the system to measure thermodynamic and kinetic parameters of protein conformational exchange on the millisecond timescale.

## Introduction

It is well established that proteins in solution are not static structures, but rather fluctuate among substates on a wide range of length and time scales, the latter typically ranging from ps-ms. The different time scales are generally correlated with the length scales, i.e., the magnitude of structural differences between states. For example, ps-ns motions of side chains and backbones are relatively low amplitude (sub-Ångstrom) fluctuations about the average structure and define many “statistical substates.” On the other hand, slow μs-ms or longer motions correspond to large-scale (multi-Ångstrom) movements of secondary structural elements, such as rigid body helix motion, or motions of entire domains that define new conformational states. Each distinct conformational state also has statistical substates. This hierarchical view of protein dynamics in solution is summarized by an energy landscape that describes the conformational free energies of the states as a function of a selected variable that represents, e.g., the root mean square deviation of atomic positions of the state relative to that of lowest energy. The funnel shaped landscape denotes the relative energies of the discrete states and the transition states that connect them (Leopold et al., 1992; Onuchic et al., 1997; Onuchic & Wolynes, 2004). The hierarchical nature of protein dynamics and the central role of protein fluctuations in evolution and function has been discussed in many reviews (Astore et al., 2024; Frauenfelder et al., n.d.; Henzler-Wildman & Kern, 2007; Mishra & Jha, 2022).

Experimental methods used to elucidate the landscape for proteins in solution include X-ray and neutron scattering as well as spectroscopic methods including nuclear magnetic resonance, Fourier-transform infrared spectroscopy, and site-directed spin labeling electron paramagnetic resonance (SDSL-EPR). SDSL-EPR is particularly attractive in that it is applicable under physiological conditions, has no restrictions as to size or complexity of the protein, and has high sensitivity for detection (picomoles-nanomoles). The capability of SDSL-EPR to site-selectively map secondary and tertiary structure, to monitor conformational transitions and protein dynamics has been extensively reviewed (Altenbach et al., 2015; Bonucci et al., 2020; Bordignon & Bleicken, 2018; Cafiso, 2014; Claxton et al., 2015; Galazzo & Bordignon, 2023; Jeschke, 2018; Roser et al., 2016; Torricella et al., 2021). The pulsed dipolar methods of double electron electron resonance (DEER)(Jeschke, 2012; Schiemann et al., 2021) and double-quantum coherence (Borbat & Freed, 1999, 2013) have greatly enhanced structure determination in proteins at cryogenic temperature, while spin relaxation enhancement methods enable inter-spin distances to be measured at ambient temperatures (Altenbach et al., 2001; Kittell et al., 2012; Rabenstein & Shin, 1995; Yang et al., 2014). Recently, variable-pressure EPR (McCoy & Hubbell, 2011) and pressure-resolved DEER (Lerch et al., 2014, 2020) have been introduced to explore conformational ensembles of proteins at equilibrium.

The inherent time scale of continuous wave (CW) X-band SDSL-EPR is on the order of ns, ideally suited to monitor side chain and fast backbone modes in spin labeled proteins. In the 0.1–100 ns time domain, motion-induced magnetic relaxation directly modulates the CW EPR spectral lineshape and information on dynamics is extracted by spectral simulations (Altenbach et al., 2015; Budil et al., 1996; Hubbell et al., 2000). Slower fluctuations in structure (conformational exchange, μs-ms) do not directly modulate the lineshape. However, the CW spectrum of a judiciously placed spin label will resolve multiple spectral components in the presence of distinct states, i.e., the spectral component corresponding to each state will not be averaged by the slow exchange event (Guo et al., 2007, 2008). In such cases, it is possible to employ pulsed saturation recovery EPR (SR-EPR) to measure the exchange rate between states on the ∼1-70 μs timescale (Bridges et al., 2010). For cases where internal motions of the spin label and overall rotational motion of the protein can be made much slower than the conformational fluctuations, the saturation transfer EPR (ST-EPR) method can extend the practical time domain to nearly 1 ms, but the conditions are relatively restrictive and the results qualitative (Beth & Hustedt, 2005). Thus, ∼1 ms is the practical upper limit for measuring exchange kinetics with SDSL-EPR using the equilibrium methods mentioned above.

To monitor large domain movements and collective motions of proteins that occur on the ms-s time scale, it is necessary to extend the time domain for detection by SDSL-EPR. To this end, perturbation-relaxation methods are promising. In these methods, a system is rapidly perturbed from an equilibrium state, and the relaxation to a new equilibrium state is followed in real time to provide exchange rate constants between the states. Of the physical perturbations possible, pressure is the most attractive for SDSL-EPR. Within the useful range of applied pressure for proteins, 0–5 kbar, the total energy of the system is not altered enough to significantly impact internal motions of side chains and backbones (Akasaka, 2006; Lerch, Yang, et al., 2015; Winter et al., 2005). Hence, any pressure-dependent EPR spectral changes are attributable to changes in protein conformation. Moreover, application of pressure to a system has the sole effect of shifting equilibria in a direction to reduce the volume of the system. Different protein conformations generally have different partial molar volumes 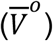, so pressure shifts a pre-existing equilibrium toward the conformation with the smallest 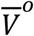 (compression effects are comparatively small in the pressure ranges used herein, ≤3500 bar) (Akasaka, 2006).

The reason that different solution conformations have different 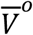 values is due to the existence of empty cavities or packing defects (Lerch, López, et al., 2015) in the “native” lowest energy state of the protein (Roche et al., 2012). Partially unfolded forms in equilibrium with the native state have a smaller 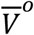 because some cavities are filled by solvent (Ando et al., 2008; Collins et al., 2005; Kamatari et al., 2011). Alternatively packed forms of the protein may also have a smaller 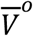 if the cavities are filled with side chains (A Vidugiris, Truckses, et al., 1970; Lerch, López, et al., 2015; López et al., 2012; Maeno et al., 2015). Both partially unfolded forms and alternatively packed forms are “excited” higher energy states of the protein that are populated by pressure. Dry molten globules may have a larger 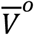 compared with the native state and would be depopulated by pressure (Neumaier & Kiefhaber, 2014).

Previous work has demonstrated the utility of variable-pressure SDSL-EPR to monitor the shift in conformational equilibria with pressure and thereby determine the corresponding volumetric and free energy differences between the states involved (Lerch et al., 2014, 2020; Lerch, López, et al., 2015; McCoy & Hubbell, 2011). In the present communication, instrumentation is introduced that allows a rapid (millisecond) jump in pressure. Immediately following a pressure jump, a protein conformational ensemble is in disequilibrium and will relax to a new equilibrium state at a rate depending on the forward and reverse rate constants for conformational exchange; this relaxation can be directly followed in real time and site-specifically by SDSL-EPR and the data analyzed to provide rate constants as well as volumetric and thermodynamic parameters of the equilibria.

Three different approaches have been used to generate changes in pressure in pressure-jump instrumentation for biological applications. One design uses a voltage-dependent extension of a piezoelectric crystal stack to drive changes in pressure (Clegg & Maxfield, 1976). A later piezoelectric design produced jumps of up to 500 bar in a time range of ∼50–100 μs (Pearson et al., 2002). These systems achieve rapid changes in pressure but are limited to jumps of relatively low magnitude due to constraints in the volume displacement that can be achieved with the actuators. The second type of system utilizes a mechanically or electrically triggered burst diaphragm to drop from high pressure to atmospheric pressure (Dumont et al., 2009). An electrically controlled burst diaphragm system was able to perform drops from 2500 bar to atmospheric pressure in ∼0.7 μs, which is an improvement in time resolution over piezoelectric designs. The inability to perform pressure jumps and drops across a range of pressures limits exploration of reaction pathways and mechanisms. Additionally, signal averaging with these systems is challenging due to the requirement to replace the diaphragm between experiments.

Here, we describe a pressure-jump system of the third type, which utilizes fast electrically controlled valves to trigger pressure jumps (Kegeles & Ke, 1975). In this system, a valve separates the sample from a reservoir, each of which are held at a different pressure. Opening of the valve triggers a rapid equilibration to an intermediate pressure; this is the pressure jump. The apparatus described here utilizes an air-operated valve system in which the flow of air is electrically controlled. The risetime of the instrument is 1–2 ms. While burst diaphragm and piezoelectric systems can achieve faster jumps, a valve system enables jumps up and down of any magnitude and to any pressure up to the maximum pressure of the system components. Computer control of the valves and pressure generator enables straightforward signal averaging and multiple measurements at different pressures to be made on a single sample.

The pressure-jump EPR system operates at Q-band (∼35 GHz). Prior high-pressure CW EPR instrumentation development has been at the more common X-band (∼9.5 GHz) frequency. In general, Q-band offers higher spin sensitivity and lower sample volumes compared with X-band. In addition to the pressure jump system, we introduce a fused silica high-pressure sample cell for Q-band that offers several advantages, including higher pressure capability, reduced background signal, and lower cost (∼$3 in materials per sample cell) compared with commercially available ceramic cells for X-band high-pressure measurements. The Q-band cells are also much simpler to fabricate compared with fused silica bundle cells for X-band (McCoy & Hubbell, 2011).

The initial application presented here utilizes the motional sensitivity of the CW lineshape to monitor a folding–unfolding equilibrium in the well-characterized model protein T4 lysozyme (T4L), illustrating the use of pressure-jump EPR to characterize the thermodynamic properties and conformational equilibria of spin-labeled proteins.

## Results

### High-pressure EPR system design and operation

The high-pressure EPR system (Figure 1) is built around an air-operated pressure intensifier and valves from Pressure BioSciences, Inc. (South Easton, MA), and is controlled on an external computer via PressureJump software written in LabVIEW (Austin, TX). The complete system includes a fused silica sample cell, loop-gap resonator, and modified Q-band microwave bridge.

**Figure 1:**
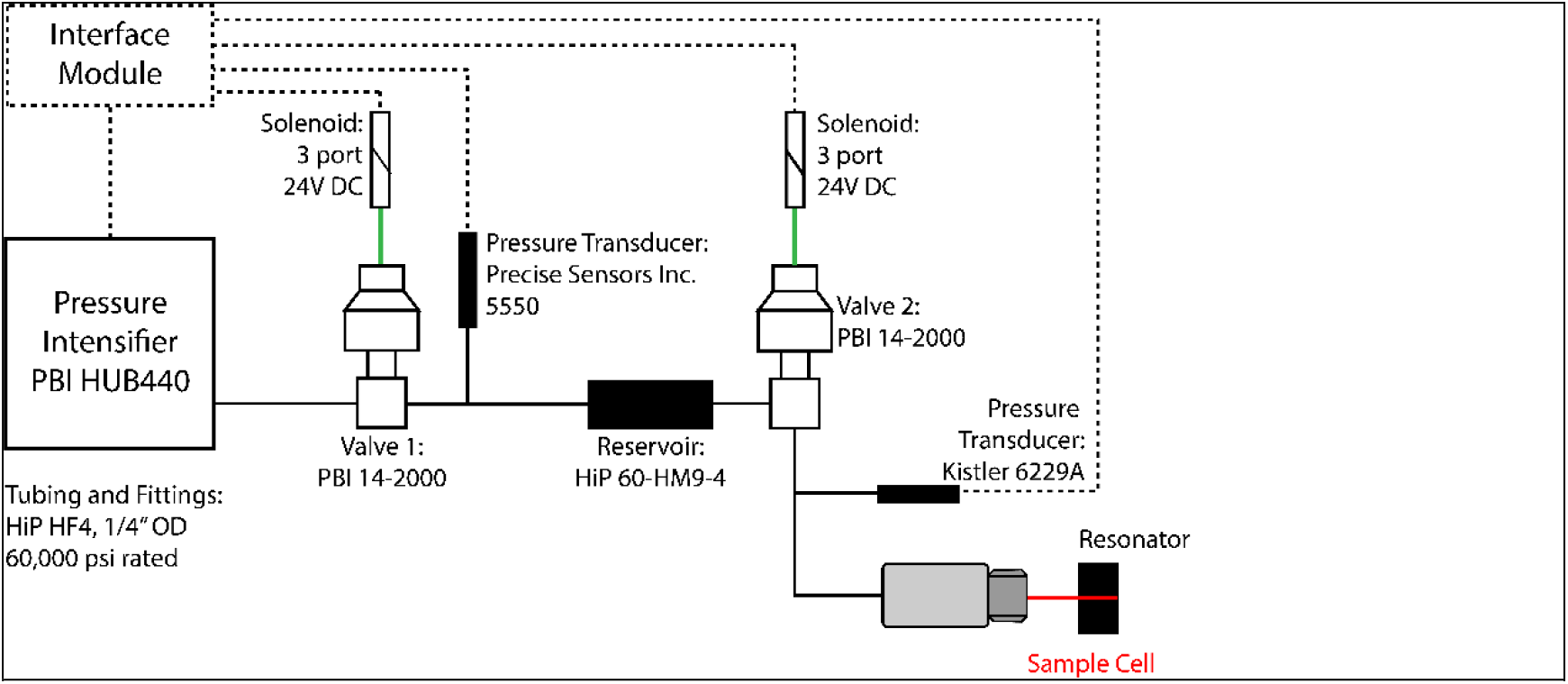
Pressure jump system design. Diagram of the pressure-jump system and its main components. Solid black connecting lines represent areas in which the pressure travels throughout the system. Dotted connecting lines indicate electrical control and data collection pathways. Green connecting lines represent compressed air lines. Red indicates the sample cell. The “Interface Module” includes the computer, the pressure-jump software interface, the DAQ I/O device, and the solid-state relays.

The Pressure BioSciences, Inc., pressure intensifier is a self-regulating system that can maintain a set pressure even in the presence of small leaks. Degassed water can be used as a pressurization liquid in this system, eliminating the need for a separator between the sample and pressurization liquid because sample diffusion is sufficiently slow (discussed below). Mechanical valve opening and closing is air operated, where the flow of air to each valve is controlled by a 3 port 24V DC solenoid. Pressure transducers from Precise Sensors, Inc. (Milford, CT), and Kistler (Winterthur, Switzerland) are used to monitor pressure within the reservoir and sample, respectively.

The high-pressure EPR system may be used for static and time-varying pressure experiments. In the static high-pressure mode of operation, both valves are open and the pressure is continuously controlled by the intensifier to maintain the set pressure. In the pressure-jump mode of operation, the software utilizes a state machine architecture with a straightforward user interface that enables control of the system pressures, timing of events, and automation of signal averaging (see Materials and Methods). To prepare the sample for a pressure-jump experiment, the sample is first pressurized to pressure P1 with both valves open, followed by closing valve 2, pressurizing the reservoir to pressure P2, and closing valve 1. The software enables variable wait times to be specified following each of these events, which allows for settling of set pressures before closing of the valves as well as equilibration of the sample at P1 prior to the jump. The pressure jump is triggered by opening of valve 2, which creates an open channel between the reservoir (10.16 cm length, 0.476 cm inner diameter, 1.81 cm^3^ volume) and sample cell (15.24 cm length, 0.02 cm inner diameter, 0.00479 cm^3^ volume), thereby allowing a rapid equilibration to a final pressure that is intermediate to P1 and P2. The pressure jump at the sample may be either a rise in pressure (P2>P1) or a drop in pressure (P2<P1).

The software records both the sample pressure and EPR signal. The data collection times prior to the jump and following the jump are adjustable settings, as is the sampling rate. To account for small variations in the timing of electronic and mechanical components of the system, the data are automatically shifted to align t=0 with the midpoint of the measured jump in pressure following opening of valve 1. The jitter in the timing of the pressure jump before alignment is typically less than 10 milliseconds (Figure S1). Following the data collection period, the system is automatically reset, and the process repeats for the user-specified number of transients.

The high-pressure sample cell (Figure 2), capable of pressures up to 6000 bar (see Materials and Methods), is manufactured from fused silica capillary tubing coated in a UV-transparent fluoropolymer sealed at one end. A copper ferrule is epoxied to the open end of the sample cell to enable connection to the pressurization system. The cell requires approximately 4 μL of sample, which is loaded by centrifugation using the assembly shown in Figure S2. The filled sample cell is connected to the outlet port on the high-pressure EPR system (Figure 1), creating a continuous column of liquid from the sample to the pressure intensifier. The fused silica pressure cells do not have a background signal, which is a distinct advantage over ceramic cells used at X-band (Lerch, Yang, et al., 2015). A small background EPR signal apparently originated from the resonator body rather than the cell (Figure S3A and B), which was easily removed by spectral subtraction (Figure S3C and D).

**Figure 2:**
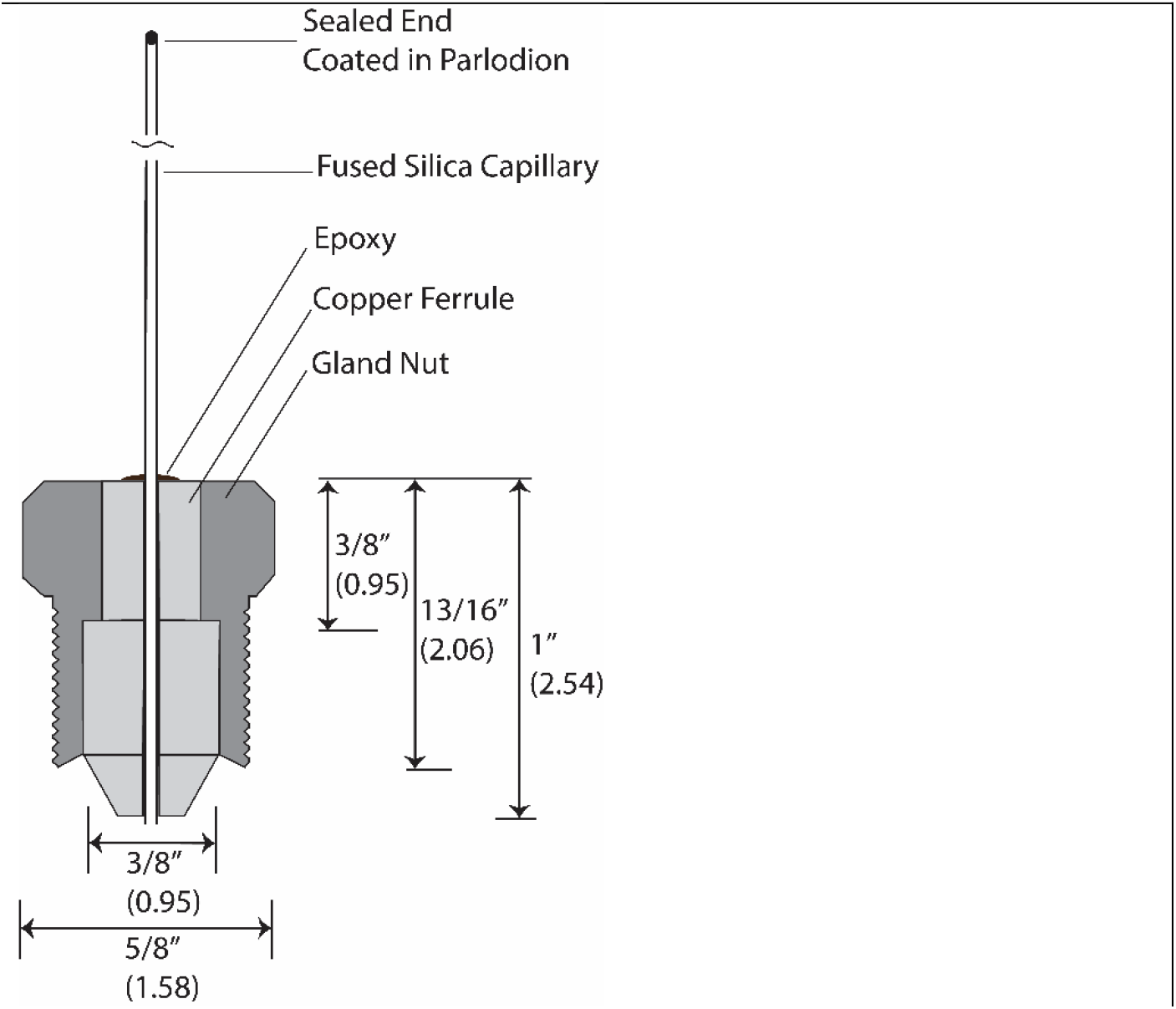
High pressure sample cell design. The gland nut and copper ferrule provide for easy connection to HF4 fittings from High Pressure Equipment Co. Dimensions here are in inches with centimeters in parenthesis.

Prior to use in a pressure-jump experiment, each pressure cell was pressure tested up to 3000 bar using the HUB440 pressure intensifier to ensure stability of each cell within the resonator. The cells are extremely robust. None of the cells that were successfully tested to 3000 bar failed during subsequent data collection, which spanned as long as 12 hours. In a separate experiment, the pressure limit of the manufactured sample cells were tested by incrementing the pressure at 1000 bar intervals up to a maximum of 6000 bar (see Materials and Methods, *HP cell materials, manufacturing, testing*). Most sample cells tested (10/15) achieved the maximum pressure of 6000 bar without failure. The HUB440 intensifier (used for high-pressure CW and pressure-jump experiments) has a maximum pressure of 4000 bar; therefore, these maximum pressure tests were performed using a HUB880 intensifier (∼7000 bar maximum).

A uniform-field three-loop–two-gap loop-gap resonator (LGR) operating at Q-band was designed for the fused silica pressure cell based on previous design for experiments at 94 GHz (Sidabras et al., 2017). The resonator body was manufactured from solid silver with modulation slots cut into the body (Figure 3). The 1 mm inner diameter of the bore was designed to accommodate the thick-walled pressure cell, which required significant reduction in the gap capacitance. Rexolite discs were added to both ends of the resonator body to enhance the uniformity of the B_1_ field within the sample loop. The resonator is housed within graphite to minimize microwave leakage through the modulation slots. A tuning screw with a copper pill is used for impedance matching. The housing assembly orients the resonator horizontally and includes sample guides and silicone rings at either end of the resonator body for proper alignment of the sample cell and minimizing transmission of mechanical vibrations from the pressurization system.

**Figure 3:**
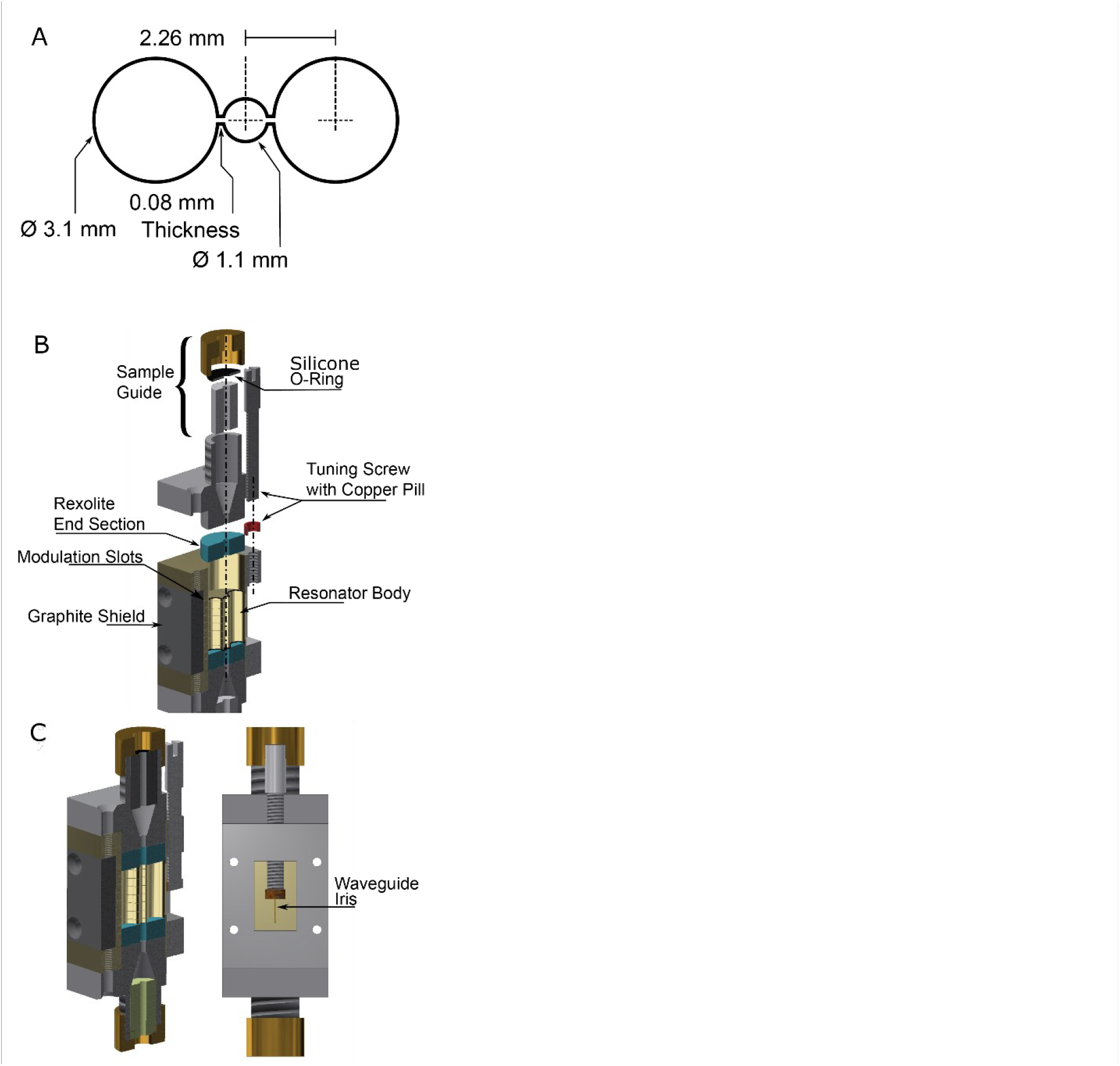
Resonator design and housing assembly (A) Cross-section view of the three-loop–two-gap 10 mm tall resonator used in this study. (B) Disassembled and (C) assembled resonator assembly highlighting the sample guiding apparatus with silicone O-ring to hold the sample and the Rexolite end sections to enhance the magnetic field uniformity over the region of interest.

A Varian E110 spectrometer bridge was modified (Figure S4) for increased stability and sensitivity. The primary modification was the addition of a low-noise amplifier to reduce phase noise at the receiver(Hubbell et al., 1987; Hyde et al., 1991). A delay line was added to the reference arm to compensate for this modification in the sample arm length. The detectors were replaced with Schottky diodes, which provide a lower noise floor. An additional arm was added to allow for automatic frequency control (AFC) lock to an external cavity for increased stability when the sample cavity is a low Q-value (<500) resonator such as an LGR; the standard high Q-value (>2000) Varian cavity (E266) was used as the lock-in cavity in our experiments.

The signal-to-noise ratio achieved by the modified spectrometer and new resonator was 446 for 100 μM TEMPOL at room temperature using spectrometer settings that are consistent with those used in variable-pressure experiments (see Materials and Methods, *Data acquisition and analysis*).

### Time scale of the physical pressure jump

The high-pressure EPR system described here can create kilobar-magnitude pressure jumps on the millisecond time scale. Multiple pressure-jump and drop speed tests ranging from approximately 500 to 2000 bar (gauge pressure) were tested (Figure 4), although jump magnitudes of up to ∼3000 bar are achievable in the current system configuration. The speed of pressure jumps is characterized herein by the time for the pressure to rise or drop from 10% to 90% of the total pressure change (Figure S5). The pressure is monitored by a transducer with a response time of 1 microsecond connected to the sample arm of the system. The opening of valve 2 initiates a rise or drop in pressure from the starting pressure to the final pressure in approximately 1–2 ms (Figure 4). This is followed by a minor (∼50 bar) decrease in pressure and small damped residual oscillations over the course of 30 milliseconds (Figures 4 and S5).

**Figure 4:**
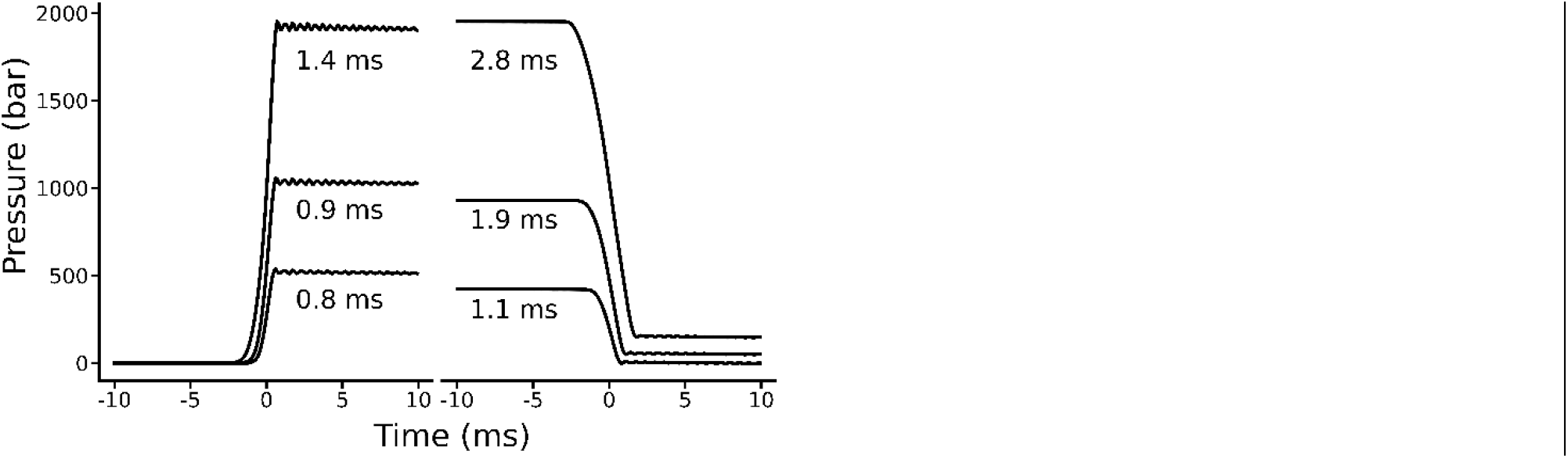
Pressure jump profile and speed. Pressure traces of pressure jumps and drops of different magnitudes. The 10–90% rise time is listed for each trace. Traces are aligned such that the midpoint of each jump or drop is aligned at 0 ms.

The jump magnitude is limited by the maximum operating pressure of the intensifier and valves (3500 bar) as well as the relative volumes of the reservoir and sample cell arms of the system. The timescale of the pressure jump is weakly dependent on the magnitude and direction of the pressure change; jump speed decreases for larger magnitude jumps and rises occur faster than drops in pressure (Figure 4).

The actual dead time of the system signal detection was limited by response times in the spectrometer, as discussed below after presentation of pressure-jump relaxation data.

### Variable-pressure Q-band CW EPR of T4L 118R1

The pressure dependence of the folded-denatured equilibrium (F↔D) of T4L 118R1 in urea was previously investigated using SDSL-EPR(McCoy & Hubbell, 2011), and this model system was used here both to demonstrate the application of the pressure-jump system to monitor the kinetics of a conformational change, and to estimate the effective dead time of the system.

Introduction of the spin label side chain R1 at the buried residue 118 (T4L 118R1; Fig. 5A) results in immobilization of the spin label in the folded state (Guo et al., 2007). In the presence of 2 M urea at ambient pressure the predominant conformation is F, although a minor amount of D is present at equilibrium. At X-band, the F↔D equilibrium for T4L in urea is clearly revealed by two spectral components for R1 at residue 118, one corresponding to an immobilized state *(i*) for R1 buried in the F conformation, and the other to a highly mobile *(m*) state in the D conformation where it is solvent exposed (McCoy & Hubbell, 2011). Pressurization results in a reversible increase in the denatured state population. The pressure-dependent, two-state equilibrium F↔D is a straightforward example of a conformational equilibrium modulated by pressure.

**Figure 5:**
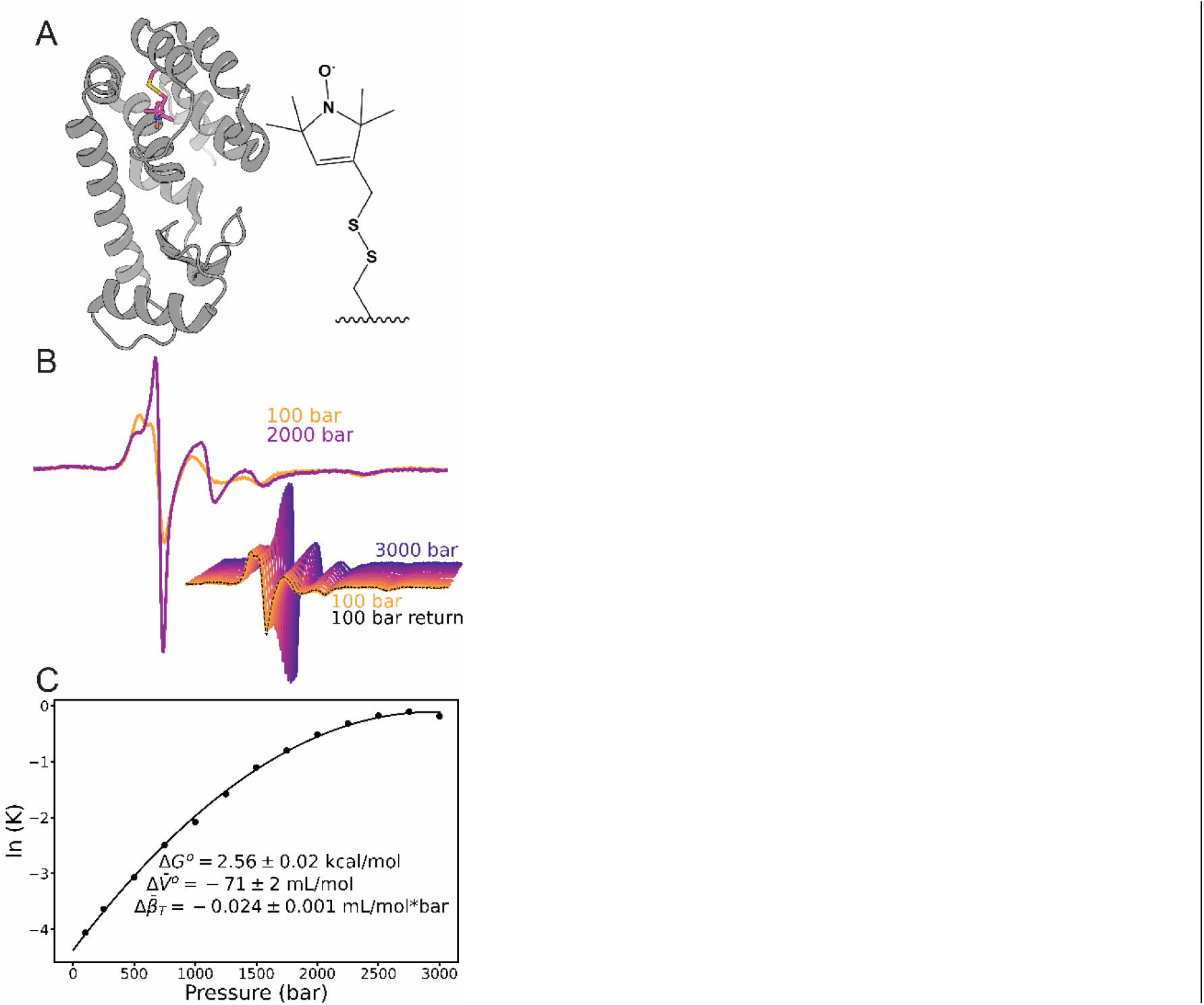
Variable-pressure CW-EPR of T4 lysozyme 118R1. (A) Crystal structure of T4 lysozyme (PDB ID: 2NTH) with the R1 side chain shown at residue 118 (magenta). Inset: structure of the R1 side chain. (B) Pressure dependence of the Q-band EPR spectra of T4L 118R1 at 300 μM; “i” and “m” denote regions of the lineshape dominated by components corresponding to immobile and mobile states, respectively. (C) The equilibrium constant determined from simulations of the variable-pressure CW EPR spectra of T4L 118R1 in 2 M urea is plotted as indicated vs. pressure. A fit to a two-state model yields the difference in free energy (Δ*G*^*o*^), change in partial molar volume 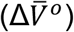, and change in compressibility 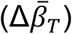 for the folded-to-denatured transition. Grey data points were not included in the fit due to inaccuracies at low pressures in this system.

The two dynamic modes *i* and *m* characteristic of distinct protein conformations were also resolved in CW Q-band spectra, as shown in Figure 5B, and increasing pressure increased the population of the D state as revealed by the increase in the *m* component.

Reversibility of pressure-populated changes was verified by collecting a spectrum at 100 bar following pressurization to 3000 bar (Figure 5B, black dotted line). The 100 bar “return” spectrum was collected after the full pressure series was completed, with static pressure and pressure jump data collected at 250 bar increments between 100 and 3000 bar. The overlay of the 100 bar spectra collected at the beginning and end of the experiment revealed negligible changes, demonstrating that the states observed at all pressures are members of the equilibrium conformational ensemble at atmospheric pressure. Additionally, the signal intensity was unchanged (data are unnormalized), confirming that sample diffusion was sufficiently slow as to negate the need for a barrier to separate sample from pressurization fluid.

### Simulation of variable pressure Q-band EPR spectra and estimation of thermodynamic equilibrium parameters

Simulations of the two-component Q-band EPR spectra as a function of pressure can be used to obtain the population ratio and hence the apparent equilibrium constant for the F↔D equilibrium as a function of pressure. From these data Δ*G*^*o*^ and 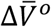 can be calculated for comparison with the values obtained from purely kinetic parameters (see below).

Simulations were performed using a simple two-state model corresponding to the F and D populations wherein the spin label in the F population had a correlation time equal to that of the isotropic rotational diffusion of the entire protein (t_R_) since it is immobilized within the folded protein interior (Lerch, Yang, et al., 2015). Because the viscosity of water increases with pressure, so does t_R_. The variation of t_R_ for T4L in the pressure range investigated here was approximately linear(McCoy & Hubbell, 2011), and this behavior is an implicit feature of the model. The spin label in the denatured state is taken to have a constant isotropic motion (t_C_) consistent with the fast motion of a disordered peptide segment; t_C_ is essentially independent of pressure within the range investigated because the dynamic unit is of effectively low molecular mass and relatively insensitive to viscosity change (McCoy & Hubbell, 2011). Thus, the simulation across the full pressure range was highly constrained with only the initial values of t_R_ and t_C_ at atmospheric pressure and the known linear pressure dependence of t_R_ required. The only variable model parameter in the simulation was then population ratio as a function of pressure. The resulting fits for this constrained model were satisfactory and shown in Figure S6. The variation of the populations determined from the simulations yielded a pressure-dependent equilibrium constant, the corresponding Gibbs free energy difference (Δ*G*^*o*^) of 2.56 ± 0.02 kcal/mol, a partial molar volume difference 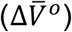 of -71 ± 2 mL/mol, and a compressibility difference 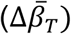 of −0.024 ± 0.001 mL/mol*bar (Figure 5C).

### Pressure jump EPR of T4 Lysozyme 118R1 in 2M urea

Example relaxation profiles following pressure jumps from 1500 bar to 2000 bar and 2500 bar to 2000 bar for T4L 118R1 in 2 M urea are shown in Figure 6. In all pressure-jump experiments reported herein, the CW EPR signal intensity at a single field position was monitored as a function of time. The signal amplitude at the minimum of the low-field line (black arrow, Figure 6 inset) was monitored in these experiments because it gives the most sensitive measure of the change in the *i*:*m* ratio, and therefore the F↔D equilibrium during pressure jump experiments. A decrease in signal at the low field line minimum indicated an increase in population of the *m* spectral component and thus a shift of the F↔D equilibrium toward the D state. The relaxation profiles in Figure 6 are consistent with a single exponential process (solid black trace).

**Figure 6:**
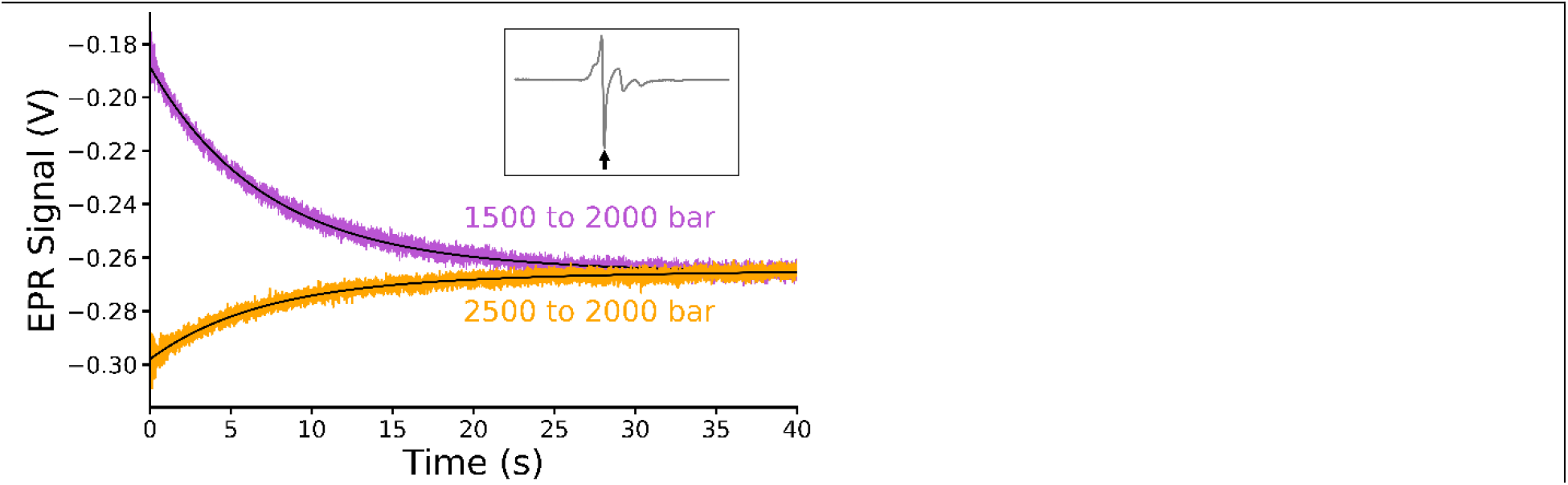
Examples of pressure-jump EPR relaxation profiles of T4L 118R1. Each relaxation profile is fitted with a one-phase decay function (black line) as part of a global fitting of the complete pressure jump dataset shown in Figure S3. Inset: EPR spectrum collected at 2000 bar. The black arrow indicates the field position monitored in pressure jump experiments.

The complete set of relaxation profiles following various pressure jumps are shown in Figure S7. The relaxation profiles were globally fit to a two-state model for conformational exchange using equation **9** (Methods) to solve for the folding and denaturing rate constants at atmospheric pressure (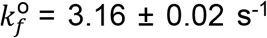 and 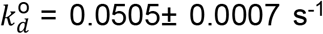, respectively) as well as the activation volumes for folding and denaturing (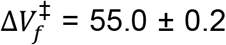 mL/mol and 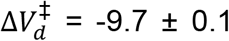 mL/mol, respectively). Table 1 gives the individual pressure-dependent time constants and rate constants derived from the fits. The natural logarithm of the exchange rate constant (1/*τ*) is plotted as a function of pressure in Figure 7, where the global fit is compared to *τ* values determined from individual fits for each relaxation profile. Increasing pressure resulted in a decrease in the rate constant up to pressures of 2500 bar, followed by a slight increase in the rate constant above 2500 bar. This Chevron-like behavior results from a pressure-dependent increase in the unfolding rate and decrease in the folding rate and is commonly observed in unfolding by chemical denaturants (Cao & Li, 2008; Jackson & Fersht, 1991; Sridevi & Udgaonkar, 2002), including unfolding of T4L by urea and guanidinium (Cellitti et al., 2007; Chen et al., 1992). The kinetic parameters from global analysis of the pressure-jump results were used to calculate a Δ*G*^*o*^ and 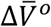 of 2.45 kcal/mol and −64.7 ± 0.2 mL/mol, respectively.

**Table 1:**
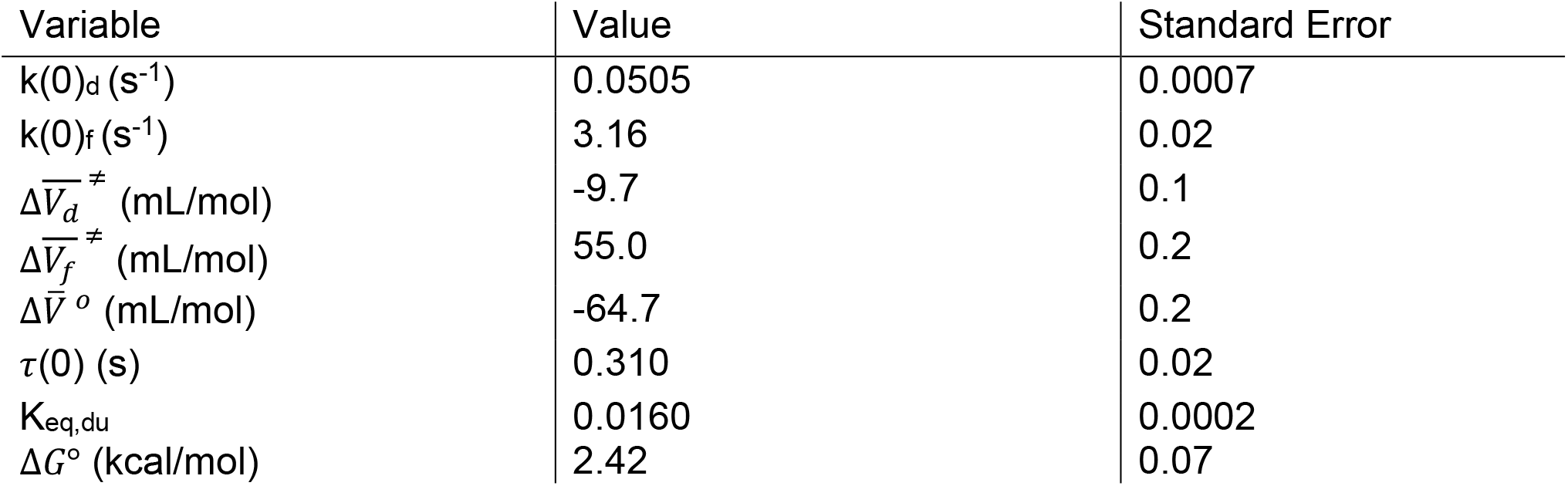
Fitted parameter values from global analysis of pressure jump relaxation profiles (Figure S3).

| Variable | Value | Standard Error |
| --- | --- | --- |
| $k(0)_d$ (s<sup>-1</sup>) | 0.0505 | 0.0007 |
| $k(0)_f$ (s<sup>-1</sup>) | 3.16 | 0.02 |
| $\Delta \bar{V}_d^\ddagger$ (mL/mol) | -9.7 | 0.1 |
| $\Delta \bar{V}_f^\ddagger$ (mL/mol) | 55.0 | 0.2 |
| $\Delta \bar{V}^o$ (mL/mol) | -64.7 | 0.2 |
| $\tau(0)$ (s) | 0.310 | 0.02 |
| $K_{eq,du}$ | 0.0160 | 0.0002 |
| $\Delta G^\circ$ (kcal/mol) | 2.42 | 0.07 |

**Figure 7:**
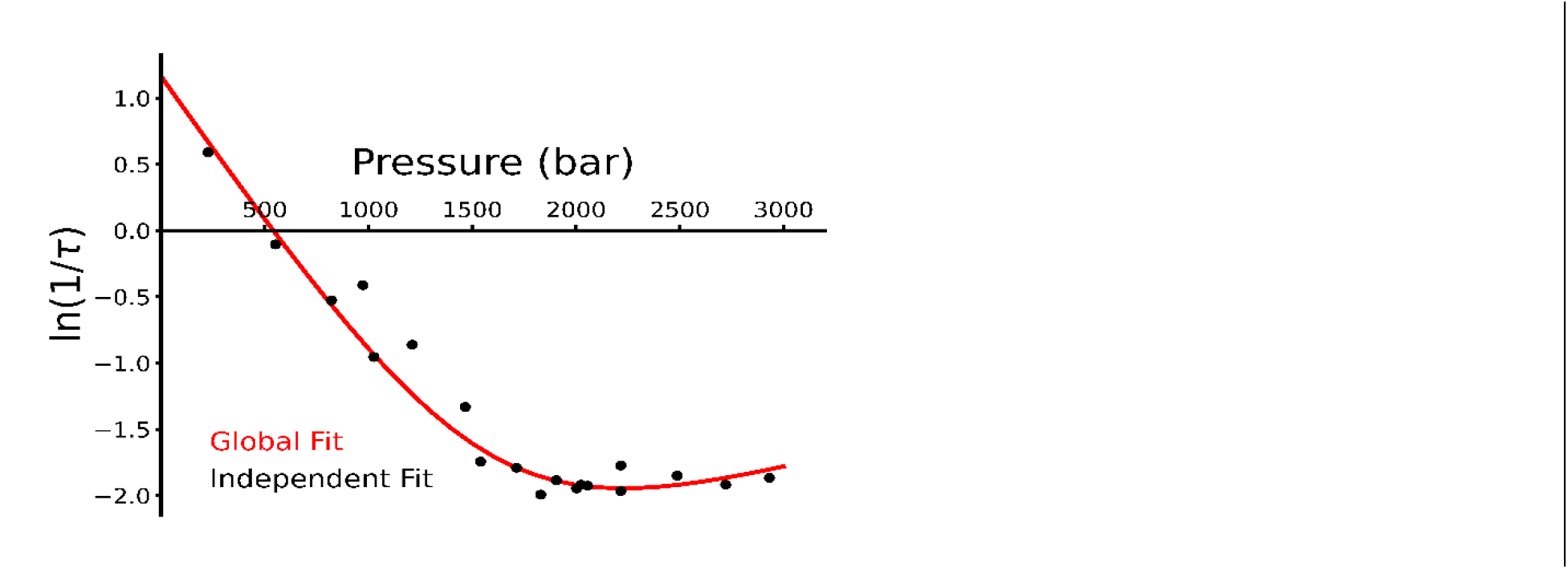
Pressure dependence of the relaxation time for F↔D exchange of T4L 118R1. The relaxation time constants determined from one phase decay fits to individual pressure jump relaxation profiles (black dots) are plotted vs. pressure. The red trace is a global fit to the data constrained by a two-state model of exchange.

### Evaluation of the system dead time using T4L118R1

Following a pressure jump, there is a rapid small magnitude increase in baseline EPR signal intensity, which then decays to a level that is very similar to the initial value. This transient was truncated from the time domain of Figure 6 but is clearly seen after the pressure jump in raw data (Figure S8, black trace). The time course of this spike in signal intensity is approximately 30 milliseconds and determines the overall deadtime of the system. A similar spike in signal was observed in both on and off resonance experiments (Figure S8B), confirming that this change in signal did not originate in the spin-labeled protein and is instead a baseline artifact. The initial 30 milliseconds of data collected following a jump are therefore excluded in the decay data analysis. The origin of this signal is apparently the response of the AFC circuit to a shift in resonance frequency that accompanies the change in water dielectric with pressure. Details are included in the supplement (Figures S9–S10).

## Materials and methods

### Protein purification and labeling

Introduction of the 118C mutation into the T4L gene (pET11a vector) was accomplished using the QuikChange site-directed mutagenesis kit (Agilent; Santa Clara, CA) and confirmed by DNA sequencing. This construct also contains the pseudo-wildtype (*WT) mutations of C54T and C97A. Expression and purification of T4L 118C was performed according to previously published methods (Fleissner et al., 2009; Sauers et al., 1992) Spin labeling of the 118C mutant was then performed as previously described (Fleissner et al., 2009). The spin labeled construct was then buffer exchanged into high-pressure buffer (20 mM MES, 2 M urea, pH 6.8) by washing 5x in a 10 kDa Amicon Ultra centrifugal filter. Spin-labeled T4L 118R1 was concentrated to 600 μM as measured by A280 (ε = 24750 cm^-1^M^-1^) and stored at 4°C for use within 24 hours. Spin label concentration was determined with a Bruker (Billerica, MA) EMXnano CW-EPR spectrometer. The ratio of spin label concentration to protein concentration yielded a labeling efficiency of 0.93 spins per protein. Protein purity was found to be >95% by SDS-PAGE

### Resonator design and assembly

A custom three-loop–two-gap resonator, illustrated in Figure 3A, was specifically designed for large access 0.665 mm outer diameter quartz tubing. The geometry was designed in Ansys Electronics Desktop (Canonsburg, PA; formerly High Frequency Structural Simulator), transferred to AutoDesk (San Francisco, CA) Inventor 3D CAD modeling software, and fabricated by Integrity Wire EDM (Sussex, WI) using electric discharge machine technology. Additionally, cross-sectional cuts were made by electric discharge machine into the 0.999-pure silver resonator body for modulation penetration. A graphite shield was placed around the silver resonator body to affix the geometry to the waveguide and remove any microwave leakage that may cause loss of signal or increased vibrational noise. The rest of the resonator assembly was fabricated by our in-house mechanic. Structural design and 3D CAD files are available upon request.

Shown in Figure 3B is the assembly of the custom resonator. Since the length of the resonator (10 mm) is beyond a free-space wavelength, it was necessary to incorporate λ_g_/4 length Rexolite end sections to improve the magnetic field distribution over the sample from sinusoidal to more uniform (Sidabras et al., 2017). A long-capacitive iris to waveguide transition was implemented to reduce non-uniform magnetic field perturbations along the sample volume and increase tuning robustness (Mett et al., 2009). Tuning was accomplished by a PTFE tuning screw with a copper pill attached to bottom. The copper pill interacts with the waveguide iris to vary the impedance match once a sample is inserted into the resonator. The sample guide with ring apparatus is designed to hold the sample in place and dampen vibrations. The sample ring (shown as black in Figure 3B) is created using a silicone disc punctured at the center for the quartz tubing to pass through. This mechanism provides adequate support for the quartz tubing while minimizing vibrations.

### High-pressure cell fabrication and testing

The high-pressure sample cell (Figure 2) consists of fused silica tubing coated in a UV-transparent fluoropolymer from Polymicro Technologies (Phoenix, AZ; part number: TSH200665) with an inner diameter of 200 μm and an outer diameter of 665 μm. Approximately 15 cm of tubing was cut with a diamond-coated file and flame sealed at one end using an oxyhydrogen torch. Due to stripping of the fluoro-polymer coating by the oxyhydrogen torch, the flame sealed end of the tubing was immediately dipped into a parlodion solution (∼3 cm^2^ parlodion, Electron Microscopy Sciences [Hatfield, PA] #19220, in 50 mL of 1:1 ethanol:diethyl ether) three times to recoat the exposed silica and keep the surface clean from any contaminants. A custom copper ferrule (Figure 3) was manufactured in-house to match the dimensions of a standard plug for ¼” high-pressure tubing from High Pressure Equipment Co. (Erie, PA; part number: 60-7HM4), except for a 0.71 mm (#70 drill bit) hole drilled through the center to accommodate the fused silica cell. The ferrule was epoxied near the open end of the sample capillary using Henkel Loctite Ablestik 24 epoxy (part number: 24 PTA) paired with Henkel Loctite Catalyst 23 LV epoxy hardener (Sycamore, IL; part number: 23LV). After curing, the empty sample cell was pressure tested to 3000 bar to avoid cell failure during EPR experiments. Pressurization fluid, either degassed water or buffer, was forced into the cell during pressurization testing, but the cell empties during depressurization such that it may be subsequently filled with sample as described above.

### Preparation of the pressure-jump EPR system and mounting the sample cell

The pressure jump system is initially primed with degassed water via four cycles of pressurization to 200 bar, bleeding through the end port, and then refilling the system with degassed water. A semi-rigid connector (Flexcoil; Pressure BioSciences, Inc.) is filled with high-pressure buffer and then attached to the loaded sample cell. Following sample cell attachment, the connector is clamped to a guide mounted at the front of the magnet and the sample cell is inserted into the resonator. The guide allows precise and controlled insertion of the sample cell into the resonator to minimize bending stress on the sample cell. The open end of the connector is then attached to the end port of the pressure jump apparatus. This creates an uninterrupted column of pressurization liquid between the pressure intensifier and the sample. Although mixing of the sample and pressurization liquid was found to be insignificant during the experiments presented herein, the use of high pressure buffer in the connector is an additional protection against mixing of the protein sample with un-buffered pressurization liquid.

### Pressure system control

Handling of system specifications between the Pressure Jump software interface (developed in LabVIEW) to the HUB440 pressure system and valves is mediated through a National Instruments (NI; Austin, TX) USB-6211 multifunction DAQ input/output (I/O) device. The NI USB-6211 is connected to a series of solid-state relays, each of which are supplied a constant voltage (24V). Relay closure by a low-power signal from the controller then connects the higher power 24V supply to a three-port 24V DC solenoid (Parker Fluid Control Division, Madison, WI). Once the electrically controlled solenoids are supplied power, air is supplied to the air-operated mechanical valves (Pressure Biosciences, Inc.; part number:14-2000) which then open, thus initiating the pressure jump. Pressure transducers from Precise Sensors, Inc. (part number: 5550) and Kistler (part number: 6229A) are used to monitor pressure within the reservoir and sample, respectively, communicated to the interface and system again through the NI USB-6211. The Pressure Jump software was used as the control interface for the pressure jump apparatus. The Pressure Jump software follows a state-machine architecture, allowing dynamic flow to other states depending on user input or programmed state progression. The software abides by the following flow of states for pressure jump experiments,

I. Pressurization of the entire system to the starting pressure (Pressure A)
II. Equilibration time (Wait A)
III. Closing of Valve 2
IV. Pressurization of the reservoir to the final pressure (Pressure B)
V. Equilibration time (Wait B)
VI. Closing of Valve 1
VII. Equilibration (Jump Delay)
VIII. Opening of Valve 2, creating the pressure jump, recording of data, and valve jitter correction (alignment of individual pressure jump traces)
IX. Fitting of accumulated data
X. I-IX repeated for “n” scans (# of scans),

where each of the above states containing a name in parentheses has specific parameters that may be set by the user in the Pressure Jump software to obtain the desired experimental specifications. This program offers simplistic automation and repeatability of pressure jump experiments as well as automatic signal averaging of collected data.

### Pressure-jump data acquisition

CW EPR was carried out at room temperature on a Varian Q-band E-110 spectrometer modified by the addition of a low-noise amplifier (Norden Millimeter, Placerville, CA; part number: N11-2927) to reduce phase noise at the receiver. Each experiment was performed using an incident power of 3.78 mW, a modulation amplitude of 1.6 Gauss, and a modulation frequency of 25 kHz. Variable-pressure CW EPR spectra were collected with a scan range of 200 Gauss, time constant of 32 ms, and sweep time of 60 s, and were repeated for nine scans. Pressure-jump relaxation profiles were collected with no time constant filter applied and repeated for 16 transients at each pressure. As described in the *Evaluation of system dead time* section, the resonant frequency is pressure dependent. As a result, the field position of the spectrum shifts as a function of pressure. Therefore, for all pressure-jump experiments on the T4L 118R1 field position were set to the low-field minimum (inset, Figure 6) at the *predetermined final pressure*. Pressure-jump data acquisition was initiated 50 ms before the pressure jump was triggered to allow accurate determination of time 0 (i.e., the time of the pressure jump). Relaxation profiles were then aligned according to the midpoint of the *pressure* transition (Figure S1) prior to signal averaging and further data analysis.

For variable-pressure and pressure-jump CW-EPR experiments, spin-labeled T4L 118R1 in high-pressure buffer was mixed in a 1:1 ratio with 50% w/w Ficoll 70 in the same buffer to give 300 μM T4L 118R1 in 25% w/w Ficoll 70 in high-pressure buffer. 4 μL of sample was loaded into the high-pressure cell by centrifugation at 4000 g for 1 minute using the assembly shown in Figure S2.

### CW spectral simulation and calculation of thermodynamic parameters for the F↔D equilibrium

Variable-pressure CW spectra were analyzed according to a two-state equilibrium between the folded and denatured protein. Variable-pressure CW EPR data were amplitude normalized and background corrected using custom software Convert&Align and Baseline, respectively (written in LabVIEW and available upon request). The pressure dependent CW EPR spectra were simulated with MultiComponent (Altenbach & Budil, 2023), which utilizes theory developed by Freed and coworkers (Budil et al., 1996) to extract dynamic parameters as well as populations to determine the apparent equilibrium constants (K) (Table S2).

The values used for the A and g magnetic tensors were g_xx_ = 2.00800, g_yy_ = 2.00557, g_zz_ = 2.00158 and A_xx_ = 7.08, A_yy_ = 6.38, A_zz_ = 35.58 for component *i*, corresponding to a nitroxide side chain in a nonpolar environment, and g_xx_ = 2.00770, g_yy_ = 2.00543, g_zz_ = 2.00235 and A_xx_ = 6.26, A_yy_ = 5.86, A_zz_ = 36.96 for component *m*, corresponding to a nitroxide side chain in an aqueous environment (Kusnetzow et al., 2006; Liang et al., 2004; Owenius et al., 2001).

As described in the text, the simulation model is defined with isotropic motion of each population with the correlation time of the immobile component increasing with pressure and that for the mobile component fixed.

Fitting parameters were optimized for each spectral component using the spectrum wherein the component was most prevalent. The spectrum of T4L 118R1 at 100 bar was used for component *i*, and the spectrum collected at 2750 bar was used for component *m*.

Least-squares fits were achieved by variation of the values of the diffusion tensor R, from which the correlation time *τ* was computed. After these values were optimized, the magnetic tensors and inhomogeneous linewidth tensor W were allowed to vary slightly to improve the fits. The diffusion tensor R of the mobile component *m* was then held constant at 1.26 ns with an inhomogeneous linewidth tensor W set to 0.71 G across the entire CW-EPR pressure range. R tensor values for component *i* increase linearly from 14.0 ns a 100 bar to 18.7 ns at 3000 bar.

The fractional population for each component of the fits (*f*_*i*_and *f*_*m*_) correspond to the population of the F and D states (*f*_*f*_ and *f*_*d*_), where *f*_*f*_ = *f*_*i*_ and *f*_*d*_ = *f*_*m*_. The equilibrium constant K for the F↔D equilibrium is given by **1**:

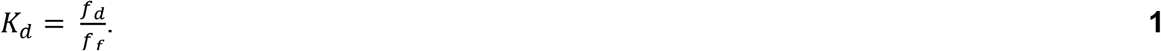

The pressure dependence of the Gibbs free energy (Δ*G*) is described by **2** and **3**:

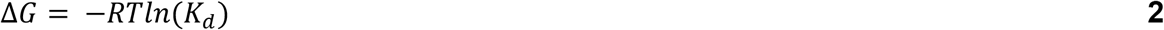

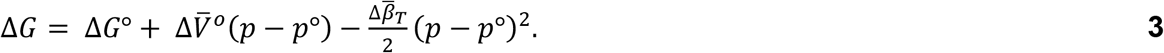

Combining equations **2** and **3** gives

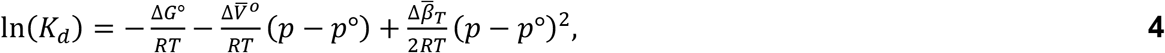

where R is the ideal gas constant and T is the temperature in kelvin. Using **4**, the pressure dependence of *K*_*d*_ obtained from simulations were used to determine Δ*G*°, 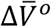, and 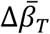.

### Analysis of P-jump relaxation data and kinetic parameters for the F↔D equilibrium

Individual relaxation profiles from pressure-jump experiments were well-fit to a single exponential decay with a linear component,

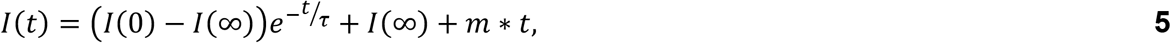

where *I*(*t*) is the EPR signal intensity at time *t, I*(0) is the intensity at time zero, *I*(∞) is the value *I* approaches asymptotically at *t* = ∞, *τ* is the relaxation time constant, and *m* is the slope of a linear component attributed to baseline drift. The baseline drift accounted for less than ∼10% of the total signal amplitude.

For a two-state reaction, *τ* at a given pressure is equal to the inverse sum of folding (*k*_*f*_(*p*)) and denaturing (*k*_*d*_(*p*)) rate constants,

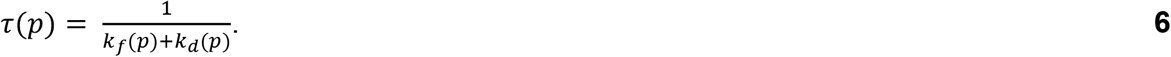

Note that p is the final pressure of a given pressure-jump experiment. According to transition state theory (Laidler & King, 1983), the rate constants are exponentially dependent on pressure through activation volumes for folding (Δ*V*^‡^_*f*_) and denaturing (Δ*V*^‡^_*d*_),

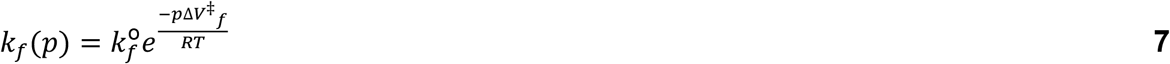

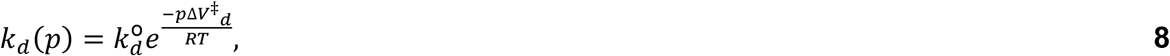

where 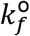 and 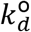 refer to the rates at zero gauge pressure (1 atm). Substituting **7** and **8** into **6** and rearranging yields:

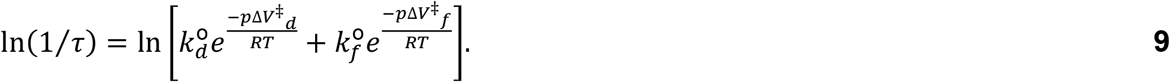

The complete set of pressure-jump relaxation profiles (Figure S3) was globally fit to **9** to determine 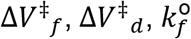, and 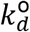. The fitted values were used to calculate the equilibrium constant and volume change of denaturation according to the following equations:

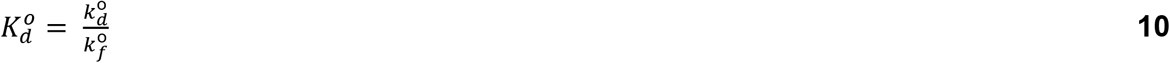

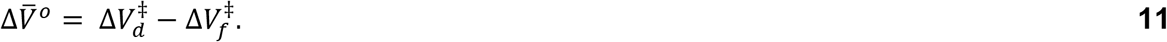

### Error analysis and comparative fitting approaches for relaxation data

Global fits to the pressure-jump data were performed using a custom LabVIEW program, the output of which provided thermodynamic parameters and error estimates based on covariance matrices. Errors were propagated through calculations as appropriate. Comparative fitting methods consisted of fitting data to later time points (i.e., excluding the first 1, 2, 4, and 6 seconds) of the relaxation profiles using one-phase decay, one-phase decay with linear component, and two-phase decay equations. Two-phase decay fits were observed having erroneous values for known parameters such as *I*(0) and *I*(∞).

Additionally, when one- and two-phase decay fits were fitted to only early timepoint data (i.e., the first 6, 8, 10 seconds of the relaxation profile), only the one-phase decay with linear component fit could be well-placed with the data in the rest of the relaxation profile. This demonstrates that a single component is present in the relaxation profile for the magnitude of jumps accessed here; therefore, a one-phase decay with a linear component is sufficient and necessary to fit these data and was incorporated into the global fitting routine. A similar approach was taken in Font et al. (Font et al., 2006).

## Discussion

### Performance of the system

Herein we describe a complete high-pressure EPR system at Q-band that offers the ability to perform both static pressure and millisecond-timescale pressure jump experiments on spin-labeled proteins. With this system, pressure-jump experiments may be performed at any pressure from atmospheric pressure up to ∼3500 bar, while static-pressure CW EPR measurements may be performed up to 6000 bar. Additionally, this system design has the advantage of allowing reversibility of pressure-populated changes to be easily verified.

System automation enables signal averaging of multiple measurements on a single sample. The active volume of sample in the resonator is only ∼0.3 μL, although ∼4 μL are needed to fill the sample cell, a value which could be reduced even further in future development.

The deadtime of the instrument is ∼30 ms, which is primarily determined by the time constant of the AFC circuitry within the bridge and is significantly longer than the pressure-jump speed (∼1–2 ms). In principle, the time resolution of the system can be further improved through modification of the AFC circuitry to improve response time.

However, shortening the dead time of the system may not generally be necessary for investigations of protein conformational exchange involving folded states. For transitions from the native state to functional low-lying excited states, pressure is expected to slow the rate of exchange because the transition state ensemble is predicted to involve expansion (i.e., dry molten globule) to allow transition between folded or packed states (Baldwin et al., 2010; Neumaier & Kiefhaber, 2014). For example, pressurization to 2000 bar increases the relaxation time constant by over an order of magnitude for the T4L denaturation reaction characterized herein. In such cases, this would allow for collecting pressure-jump data at different holding pressures and extrapolating to atmospheric pressure to determine exchange rates.

### T4L denaturation kinetics

To explore the use of the pressure-jump system to monitor protein conformational exchange, we measured denaturation kinetics for T4L in the presence of 2 M urea. One advantage of the pressure-jump approach is the ability to measure kinetics using a variety of initial conditions for each final state, because the dependence (or lack thereof) of the kinetics on initial conditions provide useful metrics for discerning the appropriate model for the reaction pathway under investigation. Relaxation profiles were collected using multiple initial pressures for each final pressure, and the time constants were independent of the initial pressure (Table S1). Variation in the observed relaxation times would indicate a reactant and/or product concentration dependence of the kinetics (Font et al., 2006), which in this case would indicate oligomerization. The lack of dependence of relaxation times on initial pressures thus indicates the reaction is mono-molecular.

The observed relaxation rate following a pressure jump is dominated by the greater of the two microscopic rate constants (see **6**); the folded state is dominant in the low pressure “limb,” whereas the denatured state dominates the high-pressure limb. Although the T4L data showing a turnover at high pressure are sparse, this Chevron-like behavior is typical in pressure-jump experiments(Desai et al., 1999; Kitahara et al., 2002; Roche et al., 2013) and indicates that pressure slows folding and speeds up denaturation. Slowing of folding kinetics with increases in pressure is commonly observed and is primarily driven by a pressure-dependent decrease in the folding rate constant (Desai et al., 1999; Kitahara et al., 2002; Roche et al., 2013). Pressure effects on the denaturation rate constant are more variable. In contrast to the results shown here for T4L, a slowing of both folding and unfolding rate constants under pressure was observed in staphylococcal nuclease (A Vidugiris, Markley, et al., 1970; Kitahara et al., 2002; Socci et al., 1996).

The pressure-jump relaxation profiles of T4L 118R1 are well-fit globally according to a two-state model for conformational exchange, yielding an expected relaxation time constant of 310 ms at atmospheric pressure. This time constant is longer than the ∼150 ms observed relaxation time constant found by stopped-flow circular dichroism at the same urea concentration, possibly due to the destabilizing effect of the spin label at residue 118 (McCoy & Hubbell, 2011).

The activation volumes in Table 1 indicate that the transition state ensemble volume is intermediate to that of the folded and denatured states, lying closer to the folded state. This may be interpreted as the transition state ensemble being collapsed with a significant portion (∼85%) of the folded-state void volume formed relative to the denatured state. This is consistent with activation volume measurements on several proteins(A Vidugiris, Markley, et al., 1970; Mohana-Borges et al., 1999; Nter Pappenberger et al., 2000), which have revealed a dehydrated transition state ensemble relative to the denatured state.

### Kinetic folding pathway from pressure vs. chemically denatured state

In contrast to the two-state model from pressure denaturation, the kinetic folding pathway of T4L from a chemically denatured state determined by stopped-flow circular dichroism includes four distinct states: a native folded state, an unfolded state, and partially folded intermediates found on either side of the rate-limiting transition state (Cellitti et al., 2007). One possible explanation for the discrepancy in the number of states observed in the kinetic folding pathway is differences in the effect of pressure and chemical denaturants on the conformational landscape. Alternatively, the difference is more likely due to the use of a local vs. global probe (Altenbach et al., 2015; McCoy & Hubbell, 2011). Site-to-site variation in the results from local probes is an important tool for revealing heterogeneity in the conformational landscape (Roche & Royer, 2018). In this case, the label at residue 118 likely reports on folding of the C-terminal domain(Guo et al., 2007; McCoy & Hubbell, 2011) and thus does not report on the intermediates observed by far-UV circular dichroism, which monitors global secondary structure.

### Use of static pressure vs. pressure-jump experiments to determine thermodynamics of the conformational landscape

The volume change of denaturation for globular proteins is typically negative at ambient temperature(Roche & Royer, 2018) due to the presence of solvent-excluded void volume in the folded protein that is eliminated upon denaturation (Akasaka, 2006). The activation volumes determined for T4L 118R1 yield a calculated volume change of denaturation 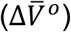 of −64.7 mL/mol, which is consistent with this model. The 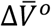 calculated from simulations of static pressure Q-band CW EPR is −71 ± 2 mL/mol. This value is in close proximity with the pressure-jump result, but differs by approximately 20 mL/mol from previous static pressure CW EPR results at X-band (McCoy & Hubbell, 2011). The difference may be due to different concentrations of the viscogen Ficoll 70 used to slow protein rotational diffusion, because molecular crowding agents, including Ficoll 70, can have significant effects on protein dynamics, kinetics, function, and folding (Kuznetsova et al., 2014; Siddiqui & Naeem, 2023; Van Den Berg et al., 1999). A key advantage of the pressure-jump approach to determine thermodynamic parameters that define the conformational landscape is that fitting of exponential decays is much more straightforward and less sensitive to user input than CW lineshape simulations.

The variable-pressure CW EPR spectra of T4L 118R1 in 2 M urea show that the spectral lineshape at Q-band (∼35 GHz) captures similar dynamic modes to X-band (McCoy & Hubbell, 2011). Traditionally, SDSL-EPR experiments using the spectral lineshape to report local protein dynamics have been performed at X-band (∼9 GHz). At X-band, the nitroxide lineshape reflects motion on the 0.1–100 ns timescale and is dominated by nitrogen hyperfine anisotropy, whereas at higher frequencies the lineshape becomes more sensitive to faster motions and spectral broadening tends to be dominated by g-anisotropy (Abergel & Palmer, 2004; Bordignon, 2017; McConnell, 1958). The m_i_ = +1 line is “compressed” from 3 mT at X-band to 2 mT at Q-band, i.e., there is less resolution of the A_z_ and A_xy_ hyperfine lines in the low-field line. Nevertheless, the two spectral components observed for T4L 118R1 in 2 M urea at X-band are easily resolvable in the Q-band spectra. Thus, the Q-band system is well-suited to investigate similar protein dynamic modes and conformational equilibria to the more traditional X-band.

## Conclusions

Taken together, the results establish the high-pressure Q-band EPR system presented here as a complete system for CW-based experiments to investigate equilibrium and kinetic properties of protein conformational ensembles under pressure. To our knowledge, this is the first high-pressure EPR system at Q-band designed for use on biological systems and the first pressure-jump EPR system of any kind. Use of an LGR at Q-band reduces the sample volume requirements compared with X-band high-pressure setups, and the fused silica cell introduced here enable higher pressures to be explored and eliminates the background signal found with ceramic pressure cells.

Sparsely populated regions of the conformational landscape may be mapped with high-pressure CW EPR methods(Lerch, López, et al., 2015; Lerch, Yang, et al., 2015; McCoy & Hubbell, 2011) as well as with pressure-resolved DEER (Lerch et al., 2014, 2020). Pressure-jump SDSL-EPR probes conformational fluctuations in the millisecond timescale, including larger-amplitude collective motions that are critical to protein function (Lange et al., 2008; Mittermaier & Kay, 2009; Yuwen et al., 2018). Although the example presented here serves to illustrate the potential of pressure-jump EPR to characterize conformational equilibria using label motion as a monitor, CW EPR enables alternative modes of detection, including solvent accessibility and distance measurements via dipolar or relaxation broadening that may be used in future studies. Additionally, pressure slows conformational exchange rates; therefore, we anticipate that this system will be useful for monitoring kinetics faster than the nominal 30 ms deadtime of the instrument. The pressure-jump method allows determination of equilibrium constants at atmospheric pressure through a straightforward and reliable exponential fit, as opposed to time consuming and often ambiguous lineshape simulations required for CW spectra. With the introduction of the system presented here, high-pressure SDSL-EPR can now be utilized to explore equilibrium structure and dynamics as well as millisecond-timescale kinetics in biomedically relevant proteins, regardless of size, complexity, or presence of a lipid bilayer.

## Acknowledgements

Research reported in this publication was supported by the National Institute of General Medical Sciences of the National Institutes of Health under award number R01GM135581 (M.L.) and the National Eye Institute under award number P30EY000311 to the Jules Stein Eye Institute. The content is solely the responsibility of the authors and does not necessarily represent the official views of the National Institutes of Health. This work was further supported by an unrestricted grant from Research to Prevent Blindness USA to the UCLA-Department of Ophthalmology and the Jules Stein Professorship Endowment to W.L.H. We are grateful for Evan Brooks’ help in expressing and purifying T4L118R1 (Hubbell Lab). We would like to thank Matt Weisbart (Stein Eye Institute, UCLA) and Richard Scherr (Biophysics Department, MCW) for their support in fabrication and assembly.

## Author Contributions

Julian D. Grosskopf: Writing – original draft; writing – review and editing; data collection; data processing; visualization.

Michael T. Lerch: Writing – original draft; writing – review and editing; pressure-jump system fabrication; conceptualization.

Wayne L. Hubbell: Writing – review and editing; conceptualization.

Christian Altenbach: Writing – review and editing; data processing; software.

Jason W. Sidabras: Writing – review and editing; resonator design & fabrication; visualization.

Jim R. Anderson: Resonator fabrication.

Richard R. Mett: Resonator design.

Robert A. Strangeway: Design of microwave bridge modification.

James J. Hyde: Resonator design.

