## Supplementary Information for "A pressure-jump EPR system to monitor millisecond conformational exchange rates of spin-labeled proteins"

Description of Supplementary Material

The Supplementary Material describes mechanical system components not shown in the main figures such as jitter correction in Figure S1 and loading of the high-pressure sample cell in Figure S2. Figure S3 describes the background signal of the resonator and high-pressure sample cell. Figure S4 highlights the modified Varian E110 spectrometer. Figure S5 demonstrates the rise time and low frequency pressure oscillations with pressure-jump experiments. Figure S6 shows static pressure continuous wave traces with corresponding MultiComponent fits. Figures S7–S9 characterize the early signal artifact of the pressure-jump system, and Figure S10 discusses the signal artifact origin. Supplementary Table 1 presents the time constants and rate constants of pressure jump experiments, and supplementary Table 2 shows the parameters used in the spectral fitting of static pressure Q band EPR data.

*System dead time is determined by response of the AFC circuit to a pressure-dependent shift in resonance frequency*

To understand the origin of this signal intensity change, it is informative to consider how the spectrometer reacts to resonator frequency changes when using an external cavity to lock the AFC (Figure S9). Initially, the signal intensity change in the first few milliseconds following a pressure jump is caused by the change in the sample resonator (LGR) resonant frequency as a result from dielectric property changes of water compression (Uematsu. M. & Franck, 1980; Yuen, 1990). This occurs both in the standard setup and external cavity setup. However, since the external cavity is unperturbed by changes during the experiment, the AFC remains locked to the external cavity frequency even though the frequency of the sample cavity has shifted lower (Figure S10A). The frequency offset between the spectrometer frequency (AFC lock) and LGR resonant frequency produces a baseline offset (Figures S9 and S10).

Using Ansoft Electronics Desktop (formerly High Frequency Structure Simulator; HFSS), we simulated the fabricated structure and estimated the frequency change and reflected power increase due to the dielectric property changes of water under compression. Extrapolating the measured data from Ref. (Uematsu. M. & Franck, 1980; Yuen, 1990), we can estimate that the change in the real part of the dielectric constant is at least an 11% increase as the pressure is raised from atmospheric pressure to 3.5 kbar. Simulations show that this increase in dielectric constant lowers the frequency by 7 MHz. Assuming the spectrometer frequency is constant, this results in an increase in the voltage standing wave ratio from 1.0964 (critically coupled, less than −30 dB) to 1.2063. For an incident power of 3.8 mW—as was used in these experiments—one can convert the voltage standing wave ratio to reflected power at the receiver, which yields 0.008 mW and 0.033 mW at atmospheric pressure and at 3.5 kBar, respectively. This power increase is equivalent to a voltage increase at the detector of approximately 0.1 V, well within the dynamic range of the bridge and on the order of the observed background shift (Figure S10).

Conversely, in the standard setup, the signal intensity returns to baseline as the AFC re-locks to the shifted LGR resonant frequency (Figure S10B). This produces the observed second phase of the “spike” as the AFC regains the lock on the sample resonator, returning the system to a similar baseline. In the standard setup, the deadtime of the instrument is thus primarily determined by the time constant of the AFC circuitry within the bridge, while in the external cavity lock setup the dead time of the instrument is determined by the settling time of the pressure and equilibration of the sample volume.

This reduction of system dead time is an advantage of the LGR and external AFC circuitry. The AFC circuit relies on the high Q-value of the external AFC to maintain lock, while the LGR—inherently low Q-value—has a much higher filling factor and overall sensitivity compared to using a high Q-value cavity as the sample cavity. Despite the offset, the EPR signal remains stable for the duration of the experiment with the external AFC circuitry while maintaining a fixed frequency and, therefore, resonance position.

To this point, we must also acknowledge the small jump in temperature that occurs concomitantly with the pressure change within the sample. Following the rapid, small amplitude jump in sample temperature, the sample will then re-equilibrate to the ambient temperature on the second timescale. This is clearly visible in data collected using an external cavity for AFC lock (Figure S9), where a low amplitude decay in EPR signal on the second timescale occurs due to the temperature-dependent frequency shift. This drift in EPR signal is eliminated by locking to the resonator where the change in frequency is accounted for.

Mechanical vibrations in the pressure-jump system as well as the pressure wave itself can cause vibrations in the sample cell that generate fluctuations in EPR signal intensity. This occurs because the resonant mode is dependent on the precise sample cell position within the sample loop of the resonator, and vibrations cause the sample cell to be displaced within the loop. To counteract these effects, the resonator housing includes alignment pieces to center the cell within the sample loop and silicone stabilizers to dampen vibrations.

Table S1: Kinetic parameters from global fit of pressure jump relaxation profiles of T4L 118R1. The first column lists the approximate initial and final pressure in each experiment, and the second column lists the actual final pressure reading from the pressure sensor at the sample.

| Pressure Jump Experiment (bar) | Final Pressure (bar) | Time Constant, global fit ($\tau$, s) | Rate Constant, global fit ($k_u$+$k_f$, $s^{-1}$) | Time Constant, individual fit ($\tau$, s) | Rate Constant, individual fit ($k_u$+$k_f$, $s^{-1}$) | Linear component slope (*m*) |
|---|---|---|---|---|---|---|
| 1250 to 250 | 227 | 0.513 | 1.950 | 0.554 | 1.806 | 0.000101 |
| 1500 to 500 | 552 | 1.022 | 0.978 | 1.110 | 0.901 | 0.000114 |
| 1750 to 750 | 821 | 1.752 | 0.571 | 1.694 | 0.590 | 0.000213 |
| 100 to 1000 | 973 | 2.322 | 0.431 | 1.510 | 0.662 | -0.000088 |
| 2000 to 1000 | 1026 | 2.553 | 0.392 | 2.596 | 0.385 | 0.000236 |
| 250 to 1250 | 1211 | 3.446 | 0.290 | 2.367 | 0.422 | -0.000177 |
| 500 to 1500 | 1466 | 4.812 | 0.208 | 3.787 | 0.264 | -0.000199 |
| 2500 to 1500 | 1540 | 5.194 | 0.193 | 5.720 | 0.175 | -0.000014 |
| 750 to 1750 | 1713 | 5.988 | 0.167 | 5.991 | 0.167 | -0.000353 |
| 2750 to 1750 | 1830 | 6.409 | 0.156 | 7.341 | 0.136 | 0.000206 |
| 1000 to 2000 | 1905 | 6.620 | 0.151 | 6.581 | 0.152 | -0.000155 |
| 1500 to 2000 | 2003 | 6.821 | 0.147 | 7.000 | 0.143 | -0.000079 |
| 3000 to 2000 | 2024 | 6.853 | 0.146 | 6.808 | 0.147 | 0.000065 |
| 2500 to 2000 | 2056 | 6.897 | 0.145 | 6.861 | 0.146 | 0.000062 |
| 1250 to 2250 | 2154 | 6.979 | 0.143 | 7.140 | 0.140 | -0.000035 |
| 1250 to 2250 | 2216 | 6.997 | 0.143 | 5.890 | 0.170 | -0.000317 |
| 1500 to 2500 | 2487 | 6.823 | 0.147 | 6.359 | 0.157 | -0.000117 |
| 1750 to 2750 | 2721 | 6.463 | 0.155 | 6.803 | 0.147 | -0.000083 |
| 2000 to 3000 | 2932 | 6.066 | 0.165 | 6.470 | 0.155 | -0.000051 |

Table S2: Fitting parameters for MOMD simulations.

| Parameter | $A_{xx}$ | $A_{yy}$ | $A_{zz}$ | $g_{xx}$ | $g_{yy}$ | $g_{zz}$ | τ (ns) | S | $\alpha_D$ | $\beta_D$ | $\gamma_D$ |
|---|---|---|---|---|---|---|---|---|---|---|---|
| Immobile Component | 7.08 | 6.38 | 35.58 | 2.00800 | 2.00557 | 2.00158 | 14.0-18.7 | 0 | 0° | 0° | 0° |
| Mobile Component | 6.26 | 5.86 | 36.96 | 2.00770 | 2.00543 | 2.00235 | 1.26 | 0 | 0° | 0° | 0° |

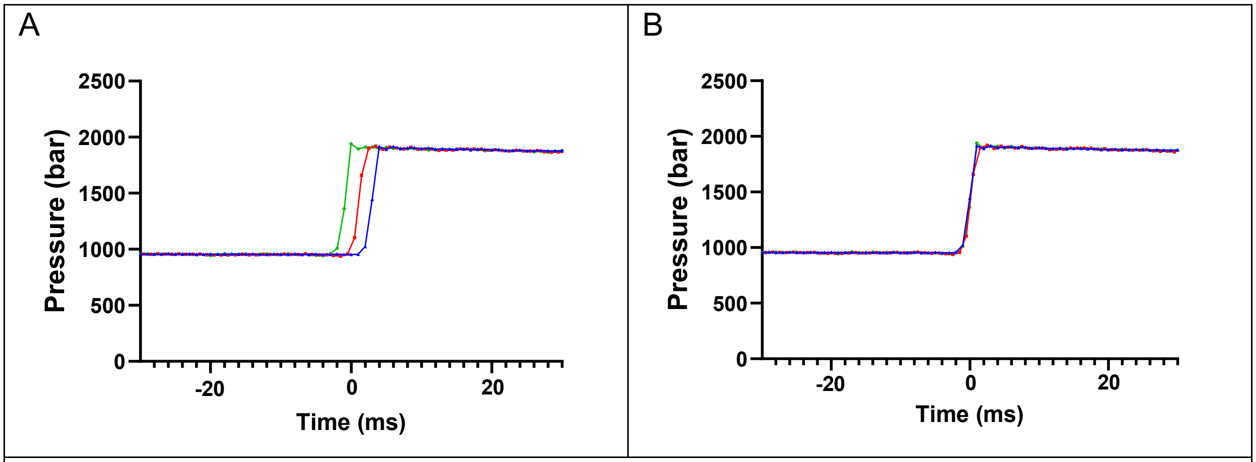


Figure S1: Pressure-jump traces before (A) and after (B) alignment via PressureJump software. Signal averaging is performed post-alignment.

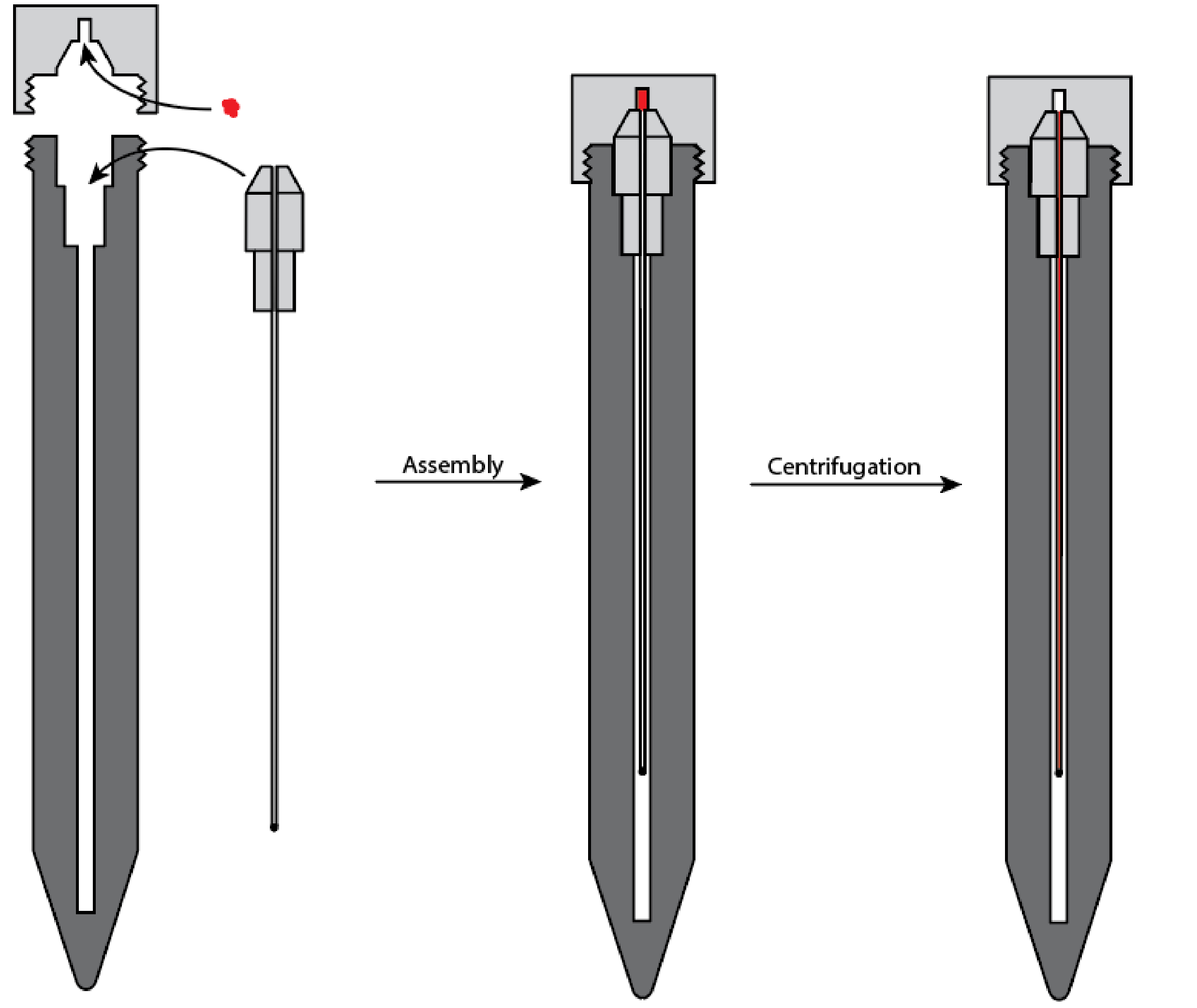


Figure S2: Apparatus and process used to load the high-pressure sample cell. A custom loader was designed and fabricated from Delrin with dimensions of a standard 15 mL Falcon tube. A 10 μL sample (red) is loaded into the reservoir in the cap (left), secured to the tube (center), and centrifuged (right).

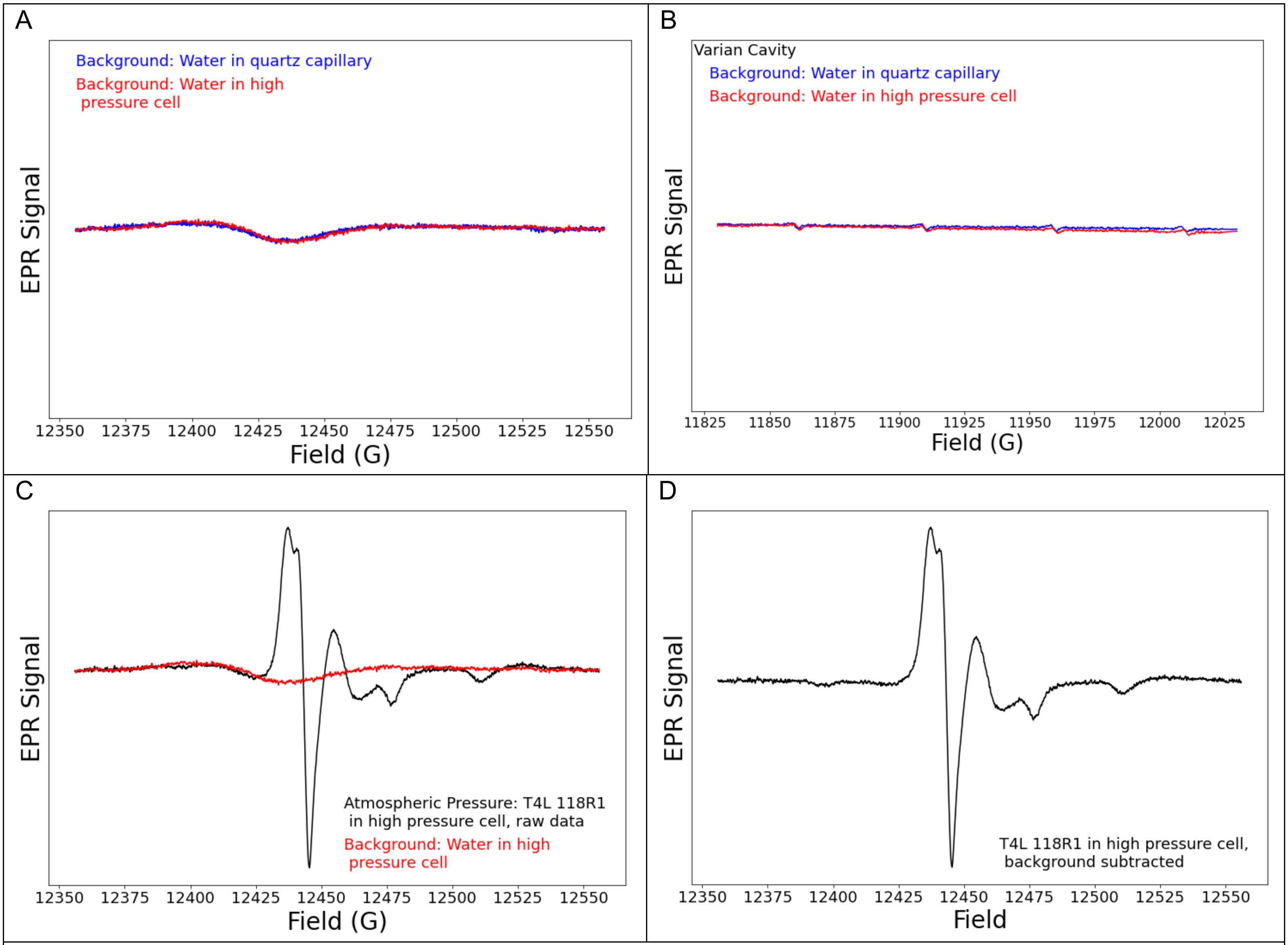


Figure S3: Background subtraction of CW EPR spectra. The background signal of water in a standard quartz capillary (blue) compared with the background signal of water in a high-pressure fused silica sample cell (red) in (A) the loop-gap resonator used for high-pressure experiments and (B) the Varian cavity. The broad, low-amplitude background signal present in the loop-gap resonator is not observed for the same samples in the Varian cavity, confirming that the background signal in A originates from the body of the resonator. Note that a small manganese impurity is present in the Varian cavity. (C) The spectrum of T4L 118R1 at atmospheric pressure (black) before subtraction of the background signal (red). (D) Background subtracted spectrum of T4L 118R1 at atmospheric pressure.

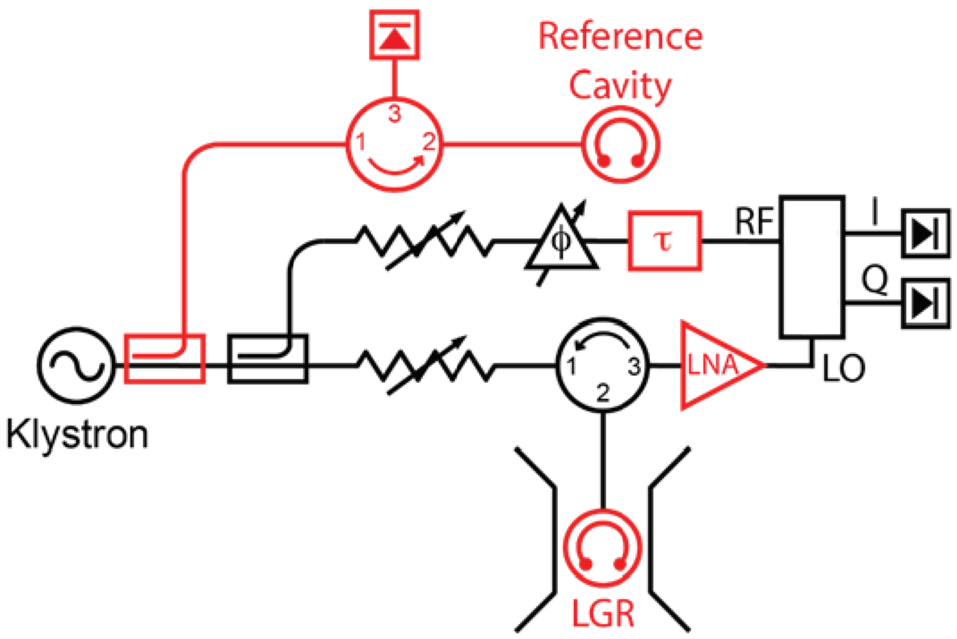


Figure S4: Schematic of the modified Varian E110 Q-band microwave spectrometer used in this work. Modifications to the spectrometer are highlighted in red. This includes: (i) The addition of a high Q-value reference cavity arm with a detector diode connected to the automatic frequency control circuit to increase stability of the circuit compared to a low Q-value sample resonator. (ii) The typical cylindrical $TE_{011}$ cavity sample resonator has been replaced with a three-loop–two-gap loop-gap resonator designed specifically for this work. (iii) The addition of a low-noise amplifier (LNA) improves the signal-to-noise-ratio at the receiver. Finally, (iv) implementation of a delay-line ($\tau$) improves phase stability and minimizes baseline changes throughout the experiment due to frequency shifts.

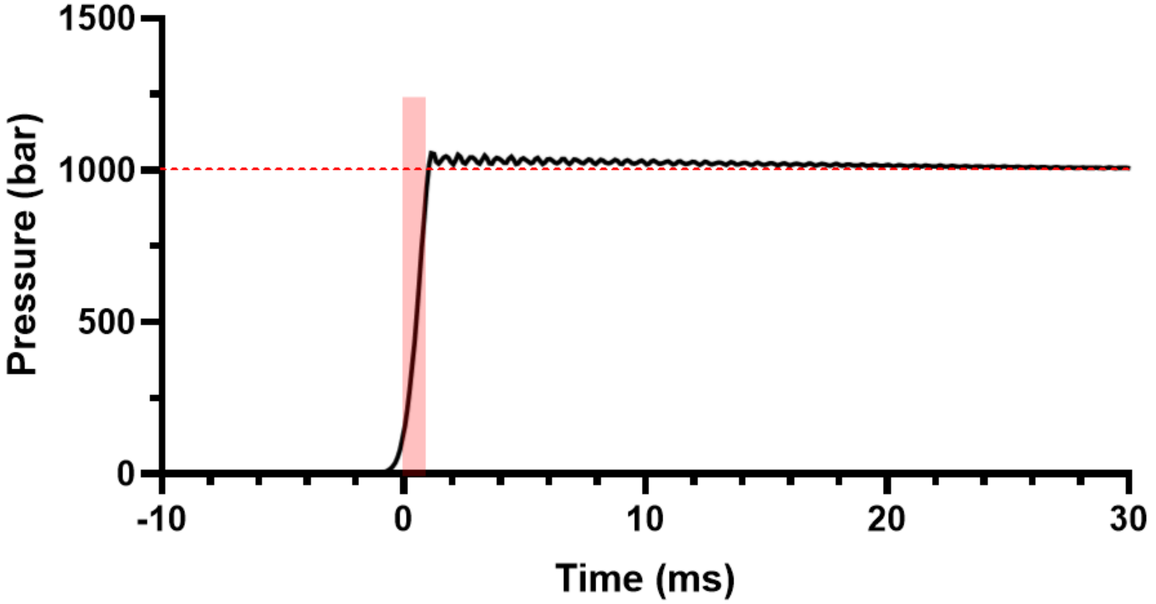


Figure S5: Rise time and low frequency oscillations in pressure-jump experiments. The 10–90% rise time is depicted by the red transparent box drawn through the jump phase of the pressure trace. The initial ~1 ms jump is followed by high frequency, low amplitude (35 bar maximum) oscillations in pressure that fully dampen out after approximately 30 ms. The red dotted line indicates the baseline pressure that is maintained in the sample cell after ~30 ms.

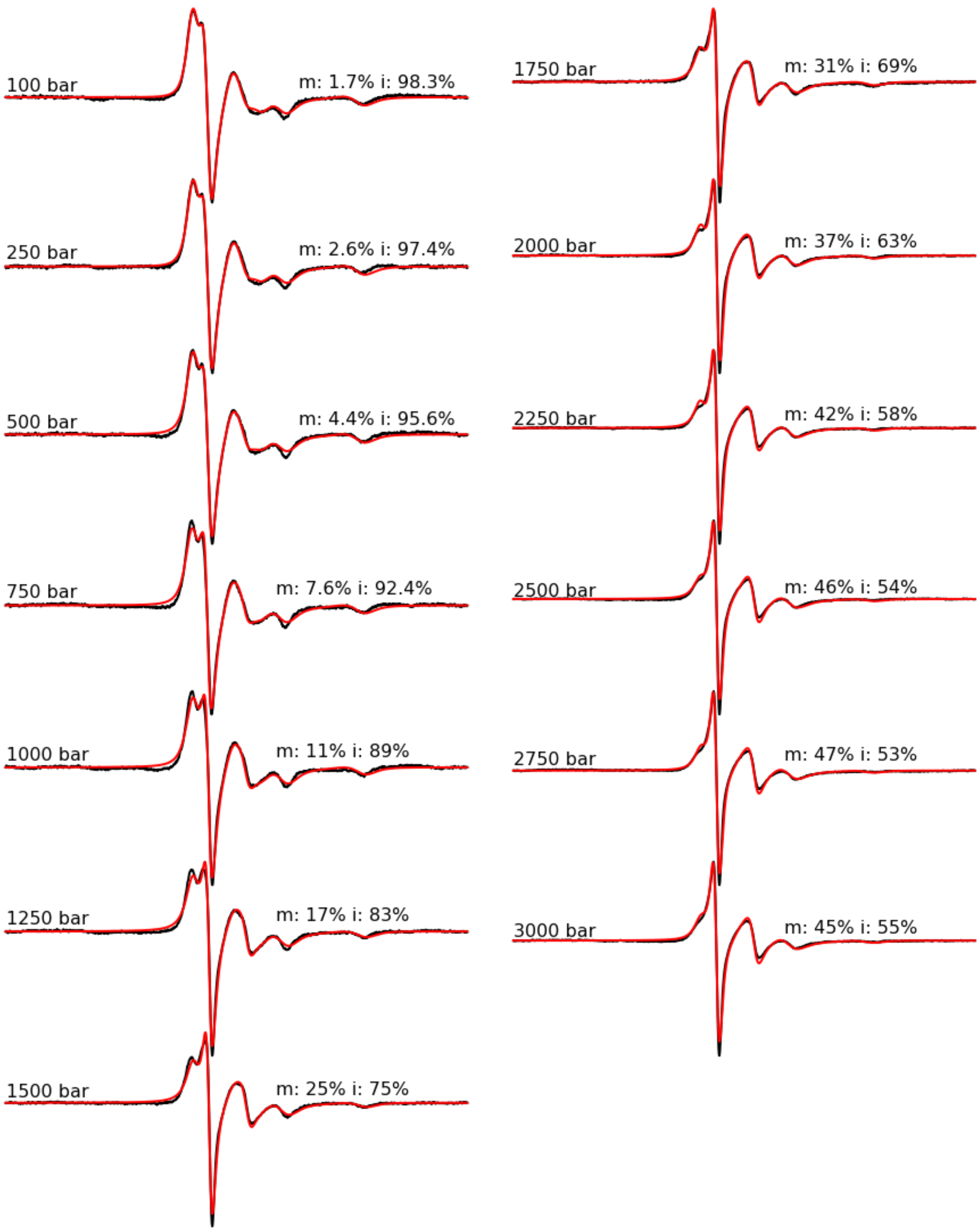


Figure S6: Experimental (black trace) and simulated (red trace) CW EPR spectra for T4L 118R1 in 2 M urea at the indicated pressure. For each, population percentages for the mobile (*m*) and immobile (*i*) components are provided.

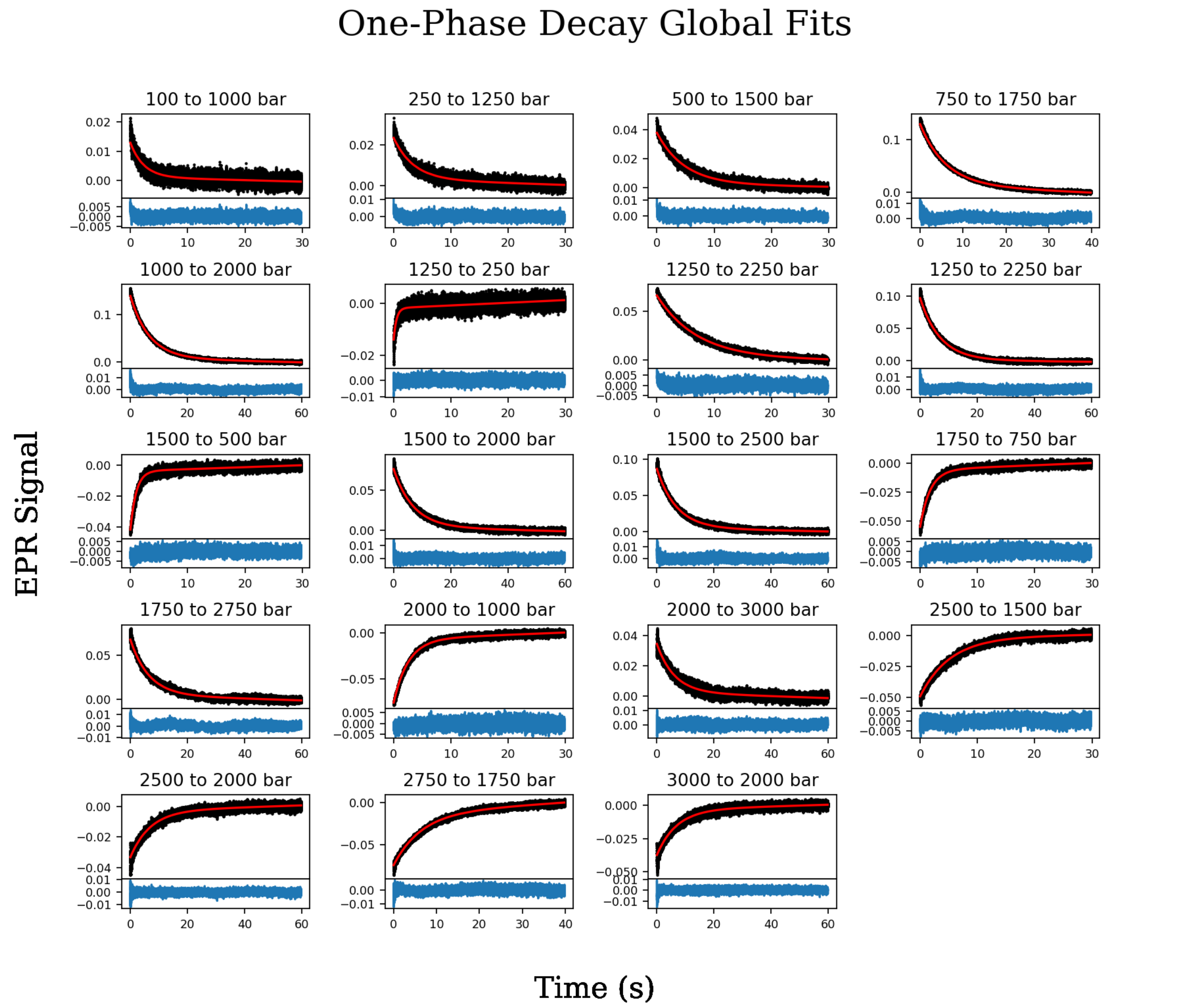


Figure S7: Complete set of pressure-jump relaxation profiles (black). Each relaxation profile is fitted with a one-phase decay function with a linear component (red) as part of a global fit to a two state model for exchange. Residuals are plotted separately below each pressure jump plot (blue).

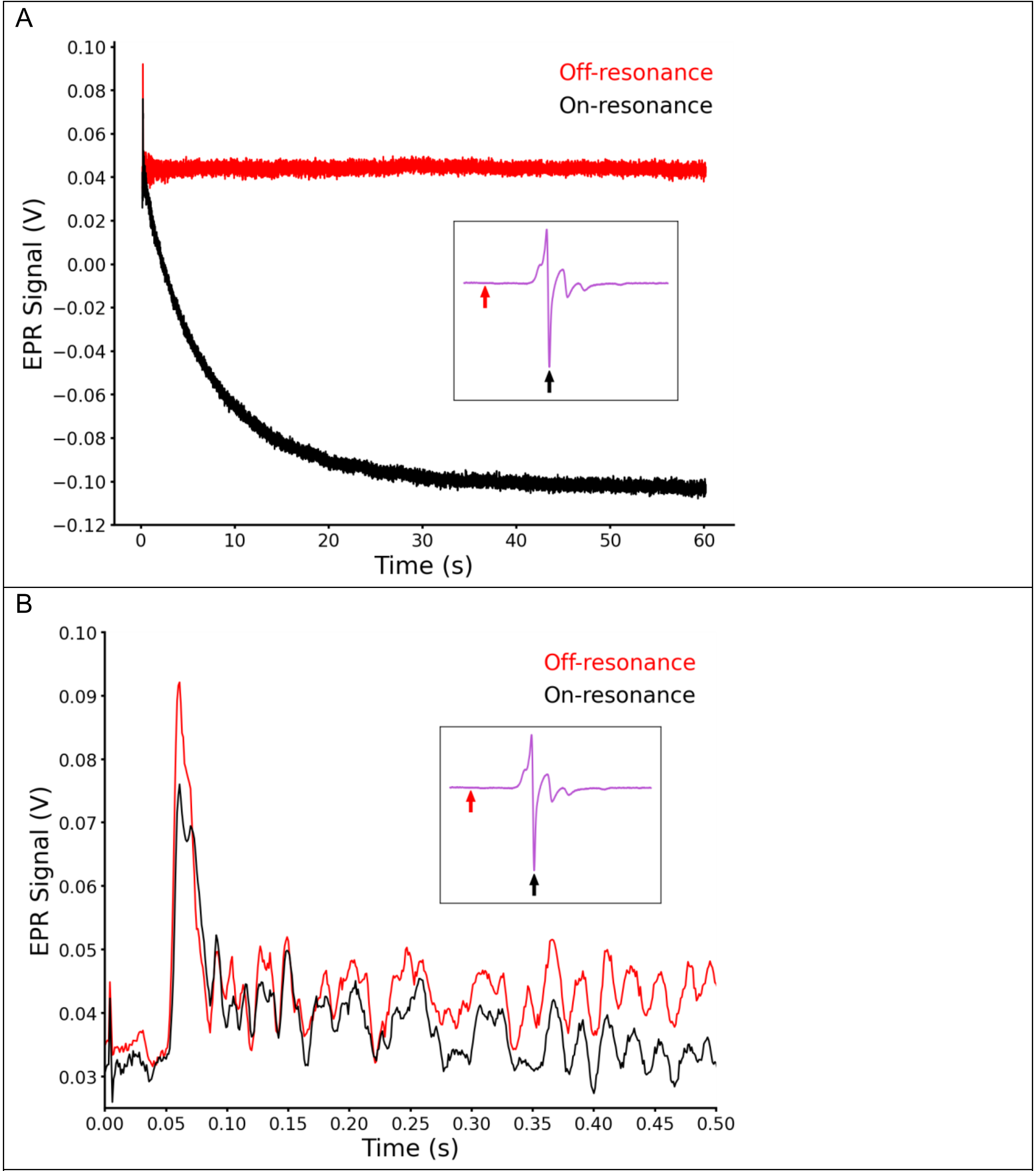


Figure S8: Comparison of signal artifact in on- vs. off-resonance pressure-jump experiments. (A) On-resonance (black) and off-resonance (red) 1000 bar to 2000 bar pressure jump EPR signal of T4L 118R1. Inset: 2000 bar spectrum with arrows marking the field position monitored for each experiment as indicated. (B) The initial 0.5 seconds of the relaxation profiles reveals a similar artifact seen in pressure jump experiments during on- and off-resonance data collection.

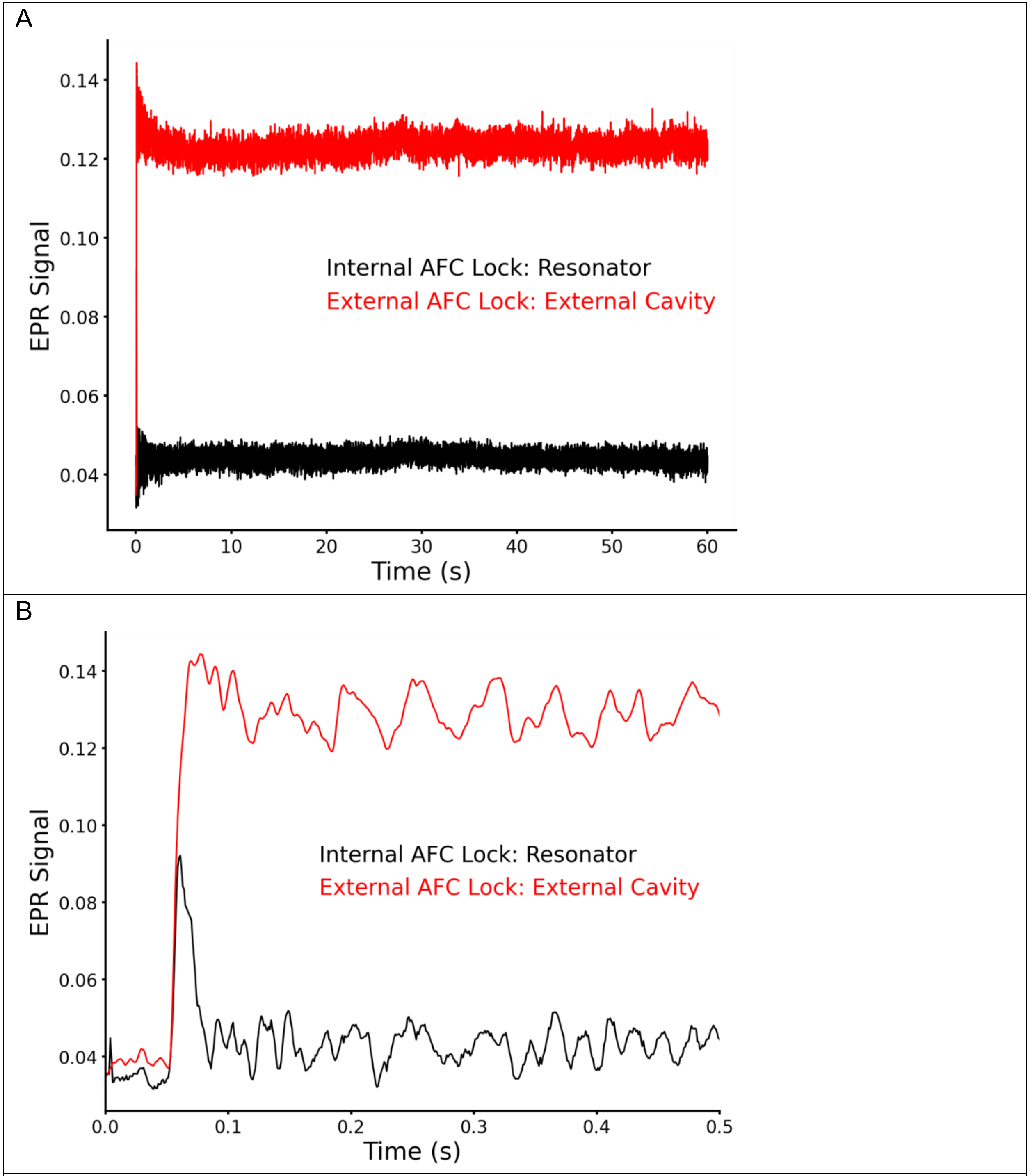


Figure S9: Comparison of signal artifact in pressure-jump experiments with AFC locked to the resonator vs. an external cavity. (A) The signal was monitored off-resonance during a pressure jump from 1000 bar to 2000 bar with AFC locked to the resonator or external cavity as indicated. (B) The initial 0.5 seconds of the trace shown in (A) reveals a difference in the early “spike” in the signal depending on the AFC lock setting.

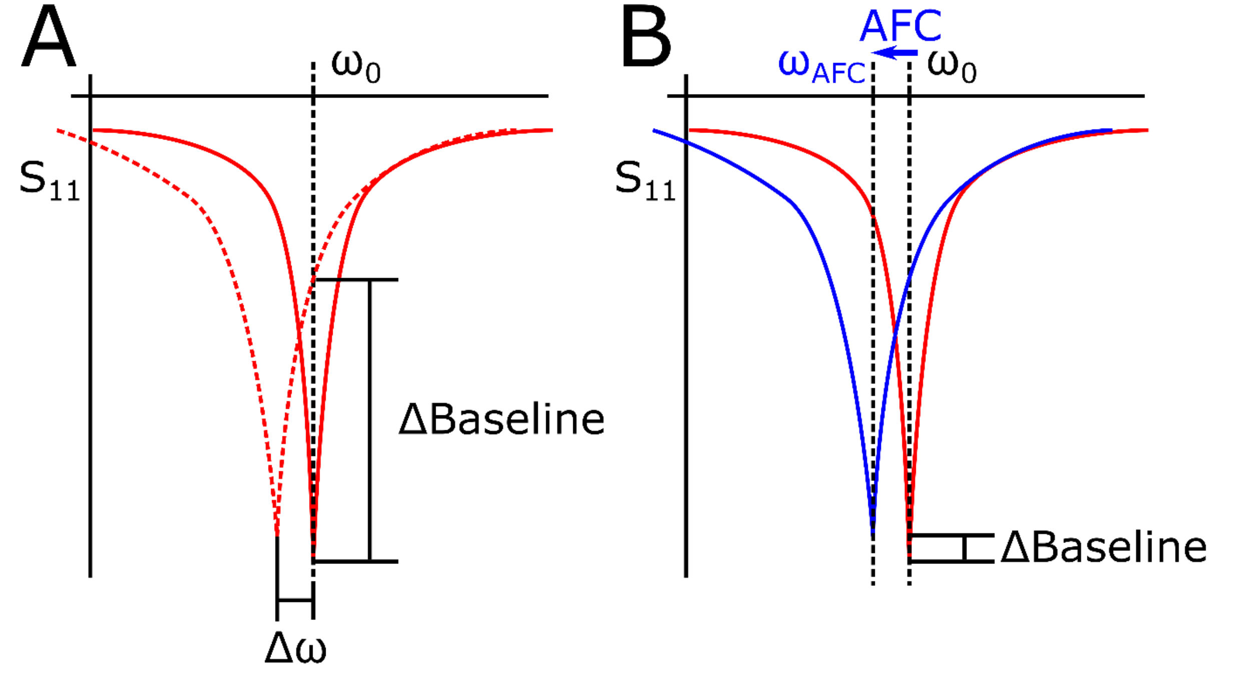


Figure S10: Diagram of the resonator profile upon rapid pressure changes and frequency correction with an external cavity AFC lock and internal cavity AFC lock. S11 represents the reflection coefficient of the resonator, the solid red line represents the resonator profile at the starting pressure, $\omega_0$ denotes the frequency at the starting pressure, $\Delta\omega$ denotes the frequency change that occurs with pressure, and ΔBaseline denotes the change in baseline signal that occurs in each case below. (A) Frequency correction using an external cavity lock. Upon pressure change, the resonator profile is changed and shifted by $\Delta\omega$ with no frequency correction (dashed red line), accounting for the change in baseline signal. (B) Frequency correction using the internal cavity AFC. Upon pressure change, the resonator profile is changed and shifted by $\omega_{AFC}$ with AFC correcting to the appropriate frequency (solid blue line), minimizing the change in baseline signal.

Uematsu. M., & Franck, E. U. (1980). Static Dielectric Constant of Water and Steam. *Journal of Physical Chemistry Reference Data*, *9*(4), 1291–1306.

Yuen, D. J. (1990). *Measurement of High Frequency Dielectric Constant and Conductivity of Fluids and Fluid-Saturated Rocks at High Pressure*. http://libraries.mit.edu/docs
